# An immunocompetent human kidney on-a-chip model to study renal inflammation and immune-mediated injury

**DOI:** 10.1101/2024.06.11.598417

**Authors:** Linda Gijzen, Marleen Bokkers, Richa Hanamsagar, Thomas Olivier, Todd P. Burton, Laura M. Tool, Mouly F. Rahman, John Lowman, Virginia Savova, Terry K. Means, Henriette L. Lanz

## Abstract

Kidney damage and dysfunction is an emerging health issue worldwide resulting in high morbidity and mortality rates. Numerous renal diseases are recognized to be driven by the immune system. Despite this recognition, the development of targeted therapies has been challenging as knowledge of the underlying mechanism and complex interactions remains insufficient. Recent advancements in the field offer promising avenues for exploring the interplay between renal cells and immune cells and their role in the development of renal inflammation and diseases. This study describes the establishment of a human immunocompetent 3D *in vitro* co-culture model of the proximal tubule in a high-throughput microfluidic platform that can be used to study renal functionality and inflammatory processes.

The model incorporated RPTEC in the top compartment and HUVECs in the bottom compartment cultured under flow and in direct contact with a collagen-I ECM gel resulting in the formation of polarized tubular structures. As an immune component, human primary monocytes of different donors were added to the lumen of the endothelium. Renal inflammation was successfully induced using complement activated serum (CAS) as evident by epithelial morphological changes, increased expression of adhesion molecules, release of pro-inflammatory cytokines, and reduced epithelial viability. Realtime migratory behavior of monocytes showed increased extravasation and migration towards the ECM and Renal compartment upon exposure to CAS with donor-to-donor differences observed. Finally, immune modulatory compounds showed efficacious inhibition of monocyte migration under inflammatory conditions in the microfluidic co-culture model.

A successful co-culture model was established and can be applied to study renal functionality in health and disease but also for drug screening due to the compatibility of the platform with automation and relatively high throughput. Overall, the described proximal tubule model has high potential to fill the gap that currently exists to study renal inflammation preclinically.

## Introduction

Kidney damage and associated dysfunction is an emerging health issue worldwide resulting in high morbidity and mortality rates.^1^ Chronic kidney disease (CKD) is a progressive irreversible condition affecting 10-15% of the population in western countries often leading to end-stage renal disease (ESRD), requiring dialysis or kidney transplantation.^2^ Over the years, treatment, and management of CKD and ESRD has evolved but despite these advances, they often address patient symptoms rather than the underlying cause of disease, with significant side effects that can further compromise quality of life.^3^ The lack of specific treatments for CKD worldwide urges the need for efficient preclinical research tools with significant predictive and translational capacity to increase our understanding of the pathophysiology of renal diseases and to support the drug development pipeline.

There are a multitude of risk factors and diseases underlying CKD with distinct pathogenic mechanisms including diabetes, hypertension, infections, (drug-induced) toxicity, and immunological disorders.^4,5^ Regardless of the different underlying mechanisms, one of the hallmarks in renal diseases is inflammation which contributes to disease development and progression.^6–8^ Inflammation is a process aiming to detect and fight harmful pathogens and subsequently promote tissue repair and recovery. However, low-grade persistent inflammation has been demonstrated as a major pathogenic factor in many renal diseases and has been recognized as both a cause and a consequence.^6^ Persistent inflammation is defined by low to moderate levels of circulating inflammation markers and the extent and effects can vary with the cause of injury. Activation of the complement system has been shown to play a crucial role in persistent inflammation.^9^ The complement system, an integral component of the innate immune system, consists of a complex network of proteins and can become activated through three separate pathways. Complement activation results in recruitment and activation of immune cells like monocytes and macrophages. Abnormalities in number and relative proportions of these cells have been described previously in CKD and specific monocyte subsets have been associated with significant loss of kidney function.^10^ These immune cells can enter the tissue and release cytokines and other signaling molecules that further drive the inflammatory response.^11^ Inappropriate activation of this tightly regulated process can contribute to cell damage, fibrosis, and declined renal function.^9,12,13^

Developing therapies that target inflammation or specifically the complement system has been of significant interest and progress has been made in identifying novel targets and developing potential therapies.^14–16^ Eculizumab, a monoclonal antibody targeting complement factor 5 (C5), showed hematological and renal improvements as well as discontinuation of dialysis in most patients with atypical hemolytic uremic syndrome (aHUS) and currently is the first-line treatment option.^17,18^ Furthermore, novel anti-inflammatory drugs targeting CCR2, IL-1β, IL-33, ASK1 and IL-6 are or have been under development for specific renal diseases.^19^ Still, most anti-inflammatory drugs fail in phase 2 clinical trials due to insufficient efficacy or increased risk of adverse outcomes such as cardiovascular events. Hence, more preclinical research is required to further expand our understanding of these complex interactions in the chronic inflammatory microenvironment allowing improved patient stratification and selection of optimal clinical endpoints, all supporting the development of anti-inflammatory therapies.

Most research in the field focuses on the proximal tubule (PT) as it is vulnerable to injury due to its high metabolic activity and exposure to high solute concentrations. PT dysfunction is linked to many acute and chronic kidney diseases.^20,21^ Numerous animal models are available to study kidney functionality and these generated valuable insights and discoveries to the field.^22,23^ Nevertheless, ethical concerns and other limitations including high costs and resources but more importantly, the ‘’translational gap’’ urges the need for alternative methods. Major advances have been made over the years in developing more sophisticated and relevant *in vitro* models of the proximal tubule. These include 3D cell culturing,^24,25^ stem cell and organoid cultures,^26–28^ 3D bioprinting,^29–31^ and organ-on-a-chip technology.^32–35^ Despite these advances, there is a demand for an *in vitro* model that integrates rapid and scalable readouts while maintaining physiological relevance. This should include features such as co-culture, fluid flow, and the absence of an artificial membrane to effectively study the proximal tubule in both health and disease.

Here, we report an immunocompetent human proximal tubule and vasculature microfluidic co-culture model to study specific aspects of kidney function and inflammation and test possible therapeutic interventions at scale. As a starting point, a previously established co-culture model was used, composed of human renal proximal tubule epithelial cells (RPTECs) and human umbilical vein endothelial cells (HUVEC) cultured as tubular structures against a collagen-I extracellular matrix (ECM) in the OrganoPlate 3-lane 40.^36^ The model was transferred to a microfluidic prototype plate, called the OrganoPlate 3-lane Narrow ECM, containing the same layout but with different channel dimensions including a smaller ECM channel. To establish an immunocompetent model, human primary monocytes were added to the lumen of the endothelial vessel. The tri-culture model revealed barrier formation, polarization, and expression of tight junction markers. Renal inflammation was induced using complement activated serum (CAS) and effects were assessed by determining marker expression, cell viability, cytokine release, and monocyte adhesion and migration. Finally, monocyte donor variability and the effect of immune modulatory compounds on monocyte migration and adhesion were assessed. Overall, we described the development of a human immunocompetent *in vitro* model of the proximal tubule that can be used to study renal functionality and inflammatory processes and has high potential to fill the gap that currently exists to study renal inflammation preclinically.

## Materials and Methods

### 2D Cell Culture

Human renal proximal tubule epithelial cells (RPTECs, SA7K clone, MTOX1030; Sigma) were cultured on PureCol-coated T75 flasks (431464U; Corning) in MEME alpha modification (M4526; Sigma) supplemented with RPTEC complete supplement (MTOXRCSUP; Sigma), L-glutamine (2.33 mM, G7513; Sigma), gentamicin (28 µg/ml, G1397; Sigma) and amphotericin B (14 ng/ml, A2942; Sigma), further referred to as RPTEC complete medium. To coat the T75 flasks, PureCol (5005-B; Advanced BioMetrix) was diluted 1:30 in cold Hank’s balanced salt solution (HBSS, H6648; Sigma) and incubated for 20 min at 37℃. The leftover solution was aspirated after which the flasks were ready. RPTECs were used for experiments up to passage 3.

Human umbilical vein endothelial cells (HUVECs, C2519A; Lonza) were cultured in T75 flasks (Nunc™ EasyFlask™, 156499; Thermo Scientific) in MV2 medium (C-22221; PromoCell) supplemented with Growth Medium MV2 SupplementMix (C-39226; PromoCell), and penicillin-streptomycin (1%, P4333; Sigma), further referred to as endothelial complete medium. HUVECs were used for experiments up to passage 5.

Cells were cultured in a humidified incubator (37℃, 5% CO_2_) and maintained by adding fresh medium every 2-3 days. Cultures were routinely tested for mycoplasma contamination and found negative.

### Monocyte isolation from human PBMCs

Human peripheral blood mononuclear cells (PBMCs) were obtained from StemExpress (PBMNC100C, 100M per vial). PBMCs were thawed, taken up in Roswell Park Memorial Institute (RPMI) 1640 basal medium (11875093; Thermo Scientific) supplemented with 10% fetal bovine serum (FBS, 16140-071; Gibco), pelleted (300g, 5 min), and washed with phosphate-buffered saline (PBS; 20012068; Sigma) containing 2% FBS. Monocytes were isolated from PBMCs by CD14 positive selection using the EasySep™ Human CD14 Positive Selection Kit (18058; Stemcell Technologies) according to manufacturer’s protocol. In short, PBMCs were pelleted (300g, 5 min) and resuspended in EasySep™ Buffer (20144; Stemcell Technologies) at a concentration of 1 x 10^8^ cells/mL and incubated with 100 µL selection antibody cocktail per mL of sample for 10 min at RT. Next, 12 µL RapidSpheres™ per mL of sample was added and incubated for an additional 3 min at RT. Cells were placed into the magnet, incubated for 10 min at RT after which the supernatant was removed and the pellet was resuspended in EasySep™ buffer. This step was repeated three times. Finally, monocytes were washed with PBS containing 2% FBS, pelleted (300g, 10 min, low brake), taken up in Cryostor® CS10 freezing medium (07930; Stemcell Technologies) at 5 x 10^6^ cells/mL and stored at -150°C.

### OrganoPlate Culture (microfluidic cell culture)

For all experiments, the OrganoPlate 3-lane 40 Narrow ECM platform was used (MIMETAS, the Netherlands, custom design). This is a custom-made microfluidic prototype plate utilizing the same chip format as the OrganoPlate 3-lane 40 platform but with different channel dimensions including a smaller ECM channel. Specifically, dimensions of the top and bottom perfusion channels are 300 µm x 165 µm (w x h), of the ECM (middle) channel 200 µm x 165 µm (w x h) and phaseguides had dimensions of 50 µm x 55 µm (w x h). The protocol for ECM loading and cell seeding was followed as previously described.^37,38^ In short, each observation window column was filled with 50 µL of Hank’s balanced salt solution (HBSS, 55037C; Sigma) per well to prevent dehydration and provide optical clarity. An extracellular matrix (ECM) composed of 4 mg/mL rat-tail collagen I (3447-020-01; AMSbio), 100 mM HEPES (15630-122; Sigma), and 3.7 mg/mL NaHCO_3_ (S5761; Sigma) was prepared and 1 µl was dispensed into the gel inlet followed by 10 min static incubation at 37℃. After polymerization of the ECM gel, 20 µL of HBSS was added to the gel inlet, and the plate was incubated overnight in a humidified incubator (37℃, 5% CO_2_). The next day, RPTECs were harvested using Accutase® solution (A6964; Sigma), pelleted (140g, 5 min) and resuspended in RPTEC complete medium at a density of 10,000 cells/µL. Subsequently, a 2 µL cell suspension was injected into the inlet of the top medium channel and placed at a 75° angle for 4.5 hours in the incubator to allow the cells to adhere to the ECM. After attachment of the cells, RPTEC complete medium was added to the remaining in- and outlet wells of the top and bottom perfusion channels and the plate was placed horizontally on an interval rocker (OrganoFlow, OFPR-L; MIMETAS, 7° inclination, 8-minute interval) in a humidified incubator (37℃, 5% CO_2_) enabling a passive, bidirectional flow through the perfusion channels.^39^ Upon flow application, the RPTECs proliferated and started lining all surfaces of the perfusion channel and formed a confluent tubule. Medium was refreshed every 2-3 days by aspirating all media from the in- and outlet wells and replacing it with fresh RPTEC complete medium.

To establish the co-culture, HUVECs were added to the bottom perfusion channel of the microfluidic chip 3 days after RPTEC seeding. HUVECs were detached using 0.025% Trypsin-EDTA (1X) solution (CC-5012; Lonza), neutralized with Trypsin Neutralizing Solution (CC-5002; Lonza), pelleted (200g, 5 min), and resuspended in endothelial complete medium at a density of 10,000 cells/µL. Media from all in- and outlet wells of the bottom perfusion channel was aspirated and 2 µL HUVEC cell suspension was injected into the inlet well. Due to loss of capillary forces by pre-wetting the channels with culture media, 1 µL of the cell suspension was taken out of the outlet well using a pipette to guide the HUVECs through the perfusion channel. This 1 µL cell suspension was added to the inlet well again and this step was repeated 2-3 times. After incubating the plate for 60 min at an angle of 75° at 37℃ for cell adherence, endothelial basal medium (CellBiologics) supplemented with growth factor supplement kit (H1168; CellBiologics) was added to the in- and outlet wells of the bottom channel. Subsequently, the media of the top channel was aspirated and replaced with fresh RPTEC complete medium. The OrganoPlate was placed back on the interval rocker in a humidified incubator (37℃; 5% CO_2_) to promote tubule formation. Polarized confluent tubules were established on day 6, at which they could be used for exposures and/or readouts. Co-cultures were used for experiments up to day 10.

### Inflammatory trigger addition and compound exposure

On day 6 of the OrganoPlate culture, the RPTEC and HUVEC tubules were exposed to media containing an inflammatory trigger and/or a potentially protective compound. To this end, Cell Biologics basal media supplemented with L-glutamine, Antibiotic-Antimycotic solution, VEGF, Heparin, EGF, FGF and hydrocortisone from the growth factor supplement kit (H1168; Cell Biologics) and 2% human AB serum (SM-612-HIS; BioIVT) was prepared and filtered using a 0.2 µm filter. This medium is further referred to as assay medium. To induce an inflammatory environment, assay medium was supplemented with 5% (v/v) cobra venom activated human complement serum (CAS, CVF-NHS; CompTech). As a negative control, assay medium was included. Media from the top channel (RPTECs) was aspirated and replaced with assay media w/ or w/o CAS (70 µL/well) and incubated for 4 hours on the interval rocker in the incubator to establish a concentration gradient in the chip. To study the effect of compounds that target either the trigger or the monocytes directly, compound A and B were tested, respectively. Both compounds were dissolved in assay medium. Compound A was added to the Renal, the Donor or to both compartments at a concentration of 25 µg/mL. Compound B was added to the Renal compartment and tested at 1, 10 and 50 µg/mL. For all compound conditions, 5% CAS (v/v) was present in the Renal compartment of the chip. After 4-hour incubation, monocytes were added to the Donor compartment, followed by addition of assay medium +/-compound A (70 µL/well). Cultures were exposed for 96 hours on a rocker platform (7° angle, 8-min interval) in the incubator (37℃; 5% CO_2_). Phase-contrast and fluorescent images were acquired every 24 hours using the ImageXpress XLS Micro High Content Imaging System (Molecular Devices).

### Monocyte cell labeling and seeding

Primary human monocytes were thawed, taken up in RPTEC 1640 medium supplemented with 10% FBS, pelleted (200g, 5min) and labelled with CellTracker Orange CMRA (C34551; Thermo Scientific) by addition of 5mL CMRA working solution. A 10 mM stock was prepared by dissolving 50 µg CMRA in 9.1 µL dimethyl sulfoxide (DMSO, D8418; Sigma). Next, a working solution of 5 µM in RPMI 1640 basal medium was prepared and warmed up to 37℃. After 30 min incubation at 37℃ in the dark, 5 mL RPMI 1640 + 10% FBS was added, cells were pelleted (200g, 5 min) and resuspended in assay medium at a density of 25,000 cells/µL.

Media from all in- and outlet wells of the bottom perfusion channel was aspirated and 2 µL monocyte cell suspension was injected into the inlet well. The same procedure as described for seeding the HUVECs was followed to guide the monocytes through the channel. Images of the first timepoint were taken, after which the plate was placed at a 75° angle for 10 min at RT to sediment the cells against the HUVEC-ECM interface. Assay medium +/- compound (70 µL/well) was added to the bottom channel and the plate was placed on a rocker platform (7° angle, 8-min interval) in the incubator (37℃; 5% CO_2_) for 96 hours.

To prevent significant monocyte loss during handling, all pipette tips and culture tubes were coated with a buffer solution. A buffer containing 10 mM Tris HCL (H5121; Promega), 0.1 mM EDTA (AM9260G; Invitrogen) and 0.1% bovine serum albumin (BSA, A2153; Sigma) in sterile Milli-Q water was prepared and used to coat all tips and tubes that were going to be in contact with monocytes. Pipette tips were dipped in the buffer to coat the outside, while the inside was coated by taken up some buffer solution using a pipette. Buffer solution was added to culture tubes, swirled around to cover the full surface, followed by aspiration of the leftover solution. Coating was performed at RT in a sterile laminar flow cabinet.

### Monocyte tracking and quantification

To track the monocytes in the chip over time, fluorescent images were taken at 0, 24, 48, 72, and 96h after addition of the cells. The images were acquired using the ImageXpress XLS Micro High Content Imaging System (Molecular Devices) at 4X magnification in combination with dichroic/emission fluorescent filters for TRITC. After imaging, the plate was placed back on the interval rocker in the incubator (37℃; 5% CO_2_). Monocyte migration was determined using Fiji (ImageJ) software as previously described.^40^ In short, a rolling ball background correction was applied to improve the signal-to-noise ratio of the acquired images. Rectangle selections were made to determine the number of monocytes in each of the following compartments separately: the Donor compartment (bottom channel; endothelium), the ECM compartment, and the Renal compartment (top channel; epithelium). Next, an automated thresholding approach was applied that created binary masks of the fluorescently labeled monocytes followed by applying a particle detection algorithm to outline and label individual monocytes. Additionally, monocytes present in the in- and outlet well of the Renal compartment were counted manually and added to the number of monocytes of the Renal compartment. Values were normalized to the number of monocytes present in the Donor compartment at the start of the assay (0h) and represented as percentages.

### Transepithelial/transendothelial electrical resistance

Transepithelial/transendothelial electrical resistance (TEER) was measured to determine the integrity and permeability of the RPTEC and HUVEC barriers in the OrganoPlate. To this end, an automated multichannel impedance spectrometer designed for use with the OrganoPlate was deployed (OrganoTEER, MI-OT-1; MIMETAS) as previously described.^41^ In short, the electrode board was cleaned with 70% ethanol (85825.360; VWR Chemicals) and was left to dry for 60 minutes in a laminar flow cabinet. Single-point baseline measurements of culture were taken on day 6. Therefore, 50 µL HBSS was added to the in- and outlet well of the ECM (middle) channel and 50 µL assay medium was added to the in- and outlets wells of the top and bottom channel. The plate was left static at room temperature (RT) for 30 minutes to equilibrate before measurement. Next, the plate was placed in the OrganoTEER device in a laminar flow cabinet and point impedance measurements were performed at RT by frequency sweep from 1000 Hz to 150 kHz (amplitude 100; precision 0.5). Data analysis was performed using the OrganoTEER software, which automatically generated the TEER values per chips in ohms (Ω). By multiplying these values with the surface area of the epithelial/endothelial ECM interface, normalized values in Ω*cm^2^ were obtained.

### WST-8 enzymatic assay

The enzymatic activity of the cultures after exposures +/-CAS was determined using the WST-8 assay (96992; Sigma). Culture medium in the OrganoPlate was replaced with 25 µL WST-8 solution diluted 1:11 in assay medium in each in- and outlet of the top and bottom perfusion channel. The plate was incubated for 30 min in the incubator (37°C, 5% CO_2_) on the rocker platform (7° angle, 8-min interval), followed by absorbance measurements at 450nm using a Multiskan FC Microplate Photometer (51119000; Thermo Scientific). As a positive control, 5% Triton X-100 (T8787; Sigma) (v/v) in assay medium was added to a few chips before addition of the WST-8 solution and incubated for 10 min on a rocker platform in the incubator. Measurements of the perfusion in- and outlets were corrected for the background signal, separately for the Renal and Donor compartment. Data were normalized to the no trigger control for either the Renal or Donor compartment and presented in percentages.

### LDH release

Supernatants from the top channel containing the RPTEC tubule and from the bottom channel containing the HUVEC vessel were collected separately after 96-hour incubation. Samples of the in-and outlet of the same chips were pooled. Lactate dehydrogenase (LDH) release was determined using the Lactate Dehydrogenase Activity Assay Kit (MAK066; Sigma) according to manufacturer’s protocol. Briefly, 2 µL sample was added in duplicate to a black glass bottom 384-well plate followed by addition of 18 µL LDH assay buffer. A NADH standard curve (20 µL/well) was added to the plate in duplicate and the plate was centrifuged (200g, 1 min) to have all solutions at the bottom of the plate. Next, 20 µL Master Reaction Mix was added to each well, placed static for 1 min at RT after which the absorbance at 450 nm was measured using the Multiskan FC Microplate Photometer (Thermo Scientific). A measurement was taken every 2 minutes for 50 minutes. Data from the timepoint before a sample resulted in a higher measurement than the values of the highest concentration of the NADH standard curve was used for analysis. Background subtraction was performed and LDH release (nmol/min per mL) was determined using the following formula:

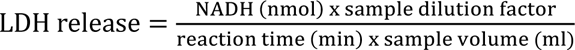

### Cytokine release

Medium samples after 96-hour incubation in the OrganoPlate were collected as previously described and used to determine the release of interleukin 6 (IL-6). Additionally, media samples of each condition prepared and used for exposure on day 6 of culture were stored (-80°C) and used for baseline measurements (referred to as 0h samples). Release was determined by using the human IL-6 DuoSet ELISA kit (DY206-05; R&D systems) in combination with ELISA Ancillary Reagent Kit 2 (DY008; R&D systems) according to manufacturer’s protocol. In short, undiluted 0-hour samples and 10x diluted 96- hour samples were added to a 96-well plate coated with the IL-6 capture antibody. Washing steps were performed using an automated plate washer (MultiFlo FX; BioTek) and absorbance was measured at 450 nm and 570 nm using the Multiskan FC Microplate Photometer (Thermo Scientific). To correct for optical imperfections, values of the 570 nm reading were subtracted from the 450 nm reading. IL-6 concentrations were calculated, corrected for the dilution factor, and presented in pg/mL.

### Immunocytochemistry

Cultures in the OrganoPlate were fixed using 3.7% formaldehyde (252549; Sigma) in PBS for 10 min, followed by two washing steps with PBS for 5 min. Subsequently, the cultures were permeabilized with 0.3% Triton X-100 (T8787; Sigma) in PBS for 10 min. Cultures were washed with 4% FBS in PBS and incubated with blocking solution (2% FBS, 2% bovine serum albumin [BSA, A2153; Sigma], and 0.1% Tween-20 [P9416; Sigma] in PBS) for 45 min. Primary antibody solution was incubated for 2,5-3h, after which the secondary antibody was incubated for 30 min. An overview of the used antibodies can be found in Table 1. Nuclei were stained with Hoechst 33342 (1:2000, H3570; Thermo Scientific) and a few chips additionally with ActinRed™ 555 reagent (R37112; Thermo Fisher). All incubation and washing steps were performed on a rocker platform (7° angle, 2-min interval) at RT. Cultures were stored in PBS and transferred to a confocal high content imaging microscope for automated imaging (Micro XLS-C, Molecular Devices). Images were acquired at 10X magnification at 2 µm increments along the height of the microfluidic channel. Image processing was performed using ImageJ to create 3D reconstructions and maximum projections.

**Table 1.**
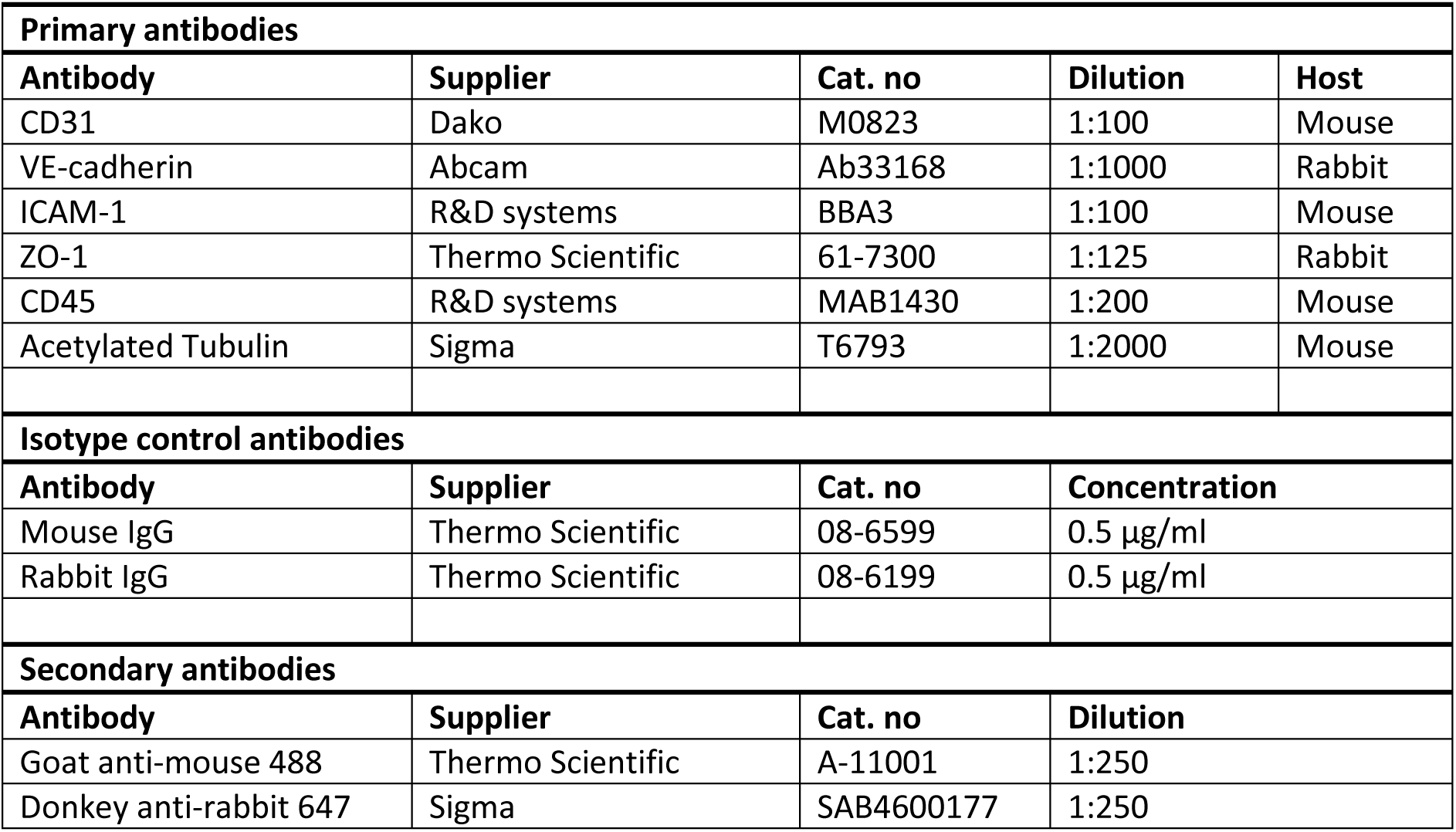
Antibodies used for immunofluorescent staining.

| Primary antibodies |  |  |  |  |
| --- | --- | --- | --- | --- |
| Antibody | Supplier | Cat. no | Dilution | Host |
| CD31 | Dako | M0823 | 1:100 | Mouse |
| VE-cadherin | Abcam | Ab33168 | 1:1000 | Rabbit |
| ICAM-1 | R&D systems | BBA3 | 1:100 | Mouse |
| ZO-1 | Thermo Scientific | 61-7300 | 1:125 | Rabbit |
| CD45 | R&D systems | MAB1430 | 1:200 | Mouse |
| Acetylated Tubulin | Sigma | T6793 | 1:2000 | Mouse |
| Isotype control antibodies |  |  |  |  |
| Antibody | Supplier | Cat. no | Concentration |  |
| Mouse IgG | Thermo Scientific | 08-6599 | 0.5 µg/ml |  |
| Rabbit IgG | Thermo Scientific | 08-6199 | 0.5 µg/ml |  |
| Secondary antibodies |  |  |  |  |
| Antibody | Supplier | Cat. no | Dilution |  |
| Goat anti-mouse 488 | Thermo Scientific | A-11001 | 1:250 |  |
| Donkey anti-rabbit 647 | Sigma | SAB4600177 | 1:250 |  |

### Statistics and Data analysis

Images were processed and analyzed using Fiji (ImageJ) software. Data analysis was performed using Excel (Microsoft Office 365; Microsoft Corp.) and GraphPad Prism (version 9.4; GraphPad Software Inc.). All data are expressed as the mean ± standard deviation (SD). Two treatment groups were compared by unpaired t-test. Multiple group comparisons were analyzed by one-way ANOVA, two-way ANOVA or mixed-effects analysis with Sidak’s or Tukey’s post hoc test. Statistical significance was indicated by one or more asterisks and considered at *p* < 0.05. Independent experiments are indicated by *N*, and replicates per experiments representing individual chips are indicated by *n*.

## Results

### Establishment of a tri-culture model of the proximal tubule

The renal proximal tubule is a critical part of the nephron responsible for blood filtration and regulating the composition of urine, which is often affected and linked to many acute and chronic renal diseases.^20^ The renal proximal tubule consists of polarized epithelial cells that display a brush border on the apical side and are supported by a basement membrane separating the tubular epithelial cells from the interstitium. To maintain the microenvironment required for proper tubule functioning, blood vessels play a crucial role and are near the epithelial cells to facilitate oxygen, nutrients, and removal of waste products. Additionally, the blood vessel can facilitate the extravasation of immune cells in response to an inflammatory stimulus as key aspect in many renal diseases (**Figure 1C**).^42^

**Figure 1.**
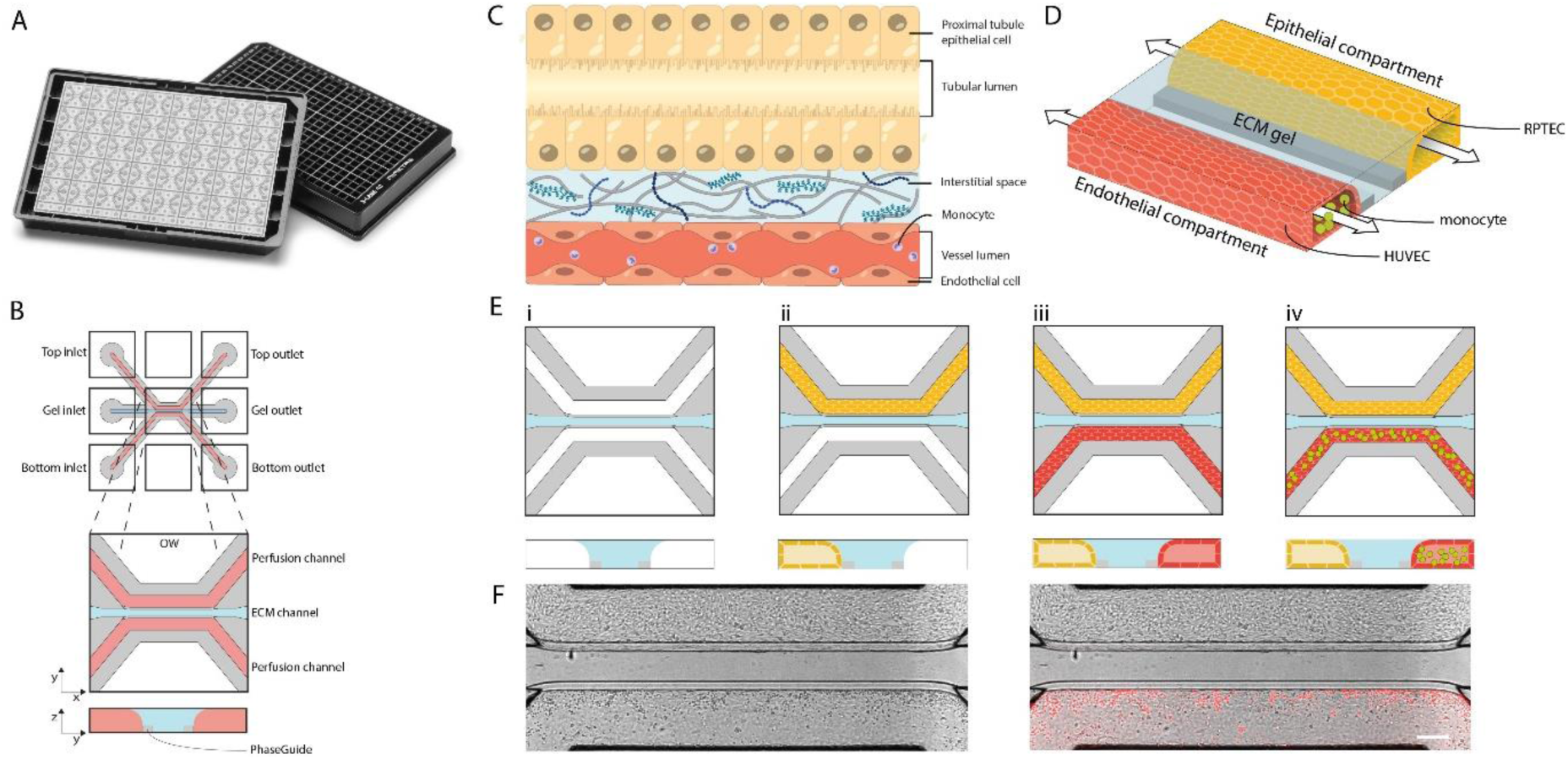
Establishment of the tri-culture kidney model in the OrganoPlate 3-lane Narrow ECM. (**A**) The OrganoPlate 3-lane, a culture platform comprised of 40 microfluidic cell culture chips embedded in a standard 384-well microtiter plate. (**B**) Schematic image of the horizontal and vertical view of one microfluidic cell culture chip consisting of 3 channels each connected to an in- and outlet well. These 3 channels join in the center of the chip (observation window; OW) and exist of 2 perfusion channels and an extracellular matrix (ECM) gel channel in the middle separated by phaseguides. (**C**) Simplistic schematic representation of the human proximal tubule composition. On top, there is a tubular structure composed of epithelial cells from the proximal tubule, featuring a lumen (yellow) through which the glomerular filtrate flows. Adjacent to the proximal tubule, there is an endothelial vessel (red) carrying blood enriched with immune cells like monocytes (purple), separated by a basement membrane composed of extracellular matrix proteins (blue). (**D**) 3D artist impression of the human tri-culture kidney model in the microfluidic chip comprising human renal proximal tubule epithelial cells (RPTEC; yellow), a collagen I ECM gel (blue), human umbilical vein endothelial cells (HUVEC; red) and monocytes (green). (**E**) Seeding strategy for establishing the tri-culture model. A collagen I ECM gel is patterned into the middle channel of the chip (i), followed by the addition of RPTECs in the top perfusion channel (ii). Next, the HUVECs are added to the bottom perfusion channel (iii) and after formation of confluent tubular structures in both channels, monocytes are added to the lumen of the endothelial vessel in medium (iv). (**F**) Phase contrast with fluorescent overlay image of the model comprising a RPTEC tubule in the top perfusion channel and a HUVEC vessel with perfused fluorescently labeled monocytes in the bottom perfusion channel at day 6 of culture, right after adding the monocytes. Scale bars in white = 200 µm.

The human proximal tubule environment was modeled using epithelial, endothelial, and immune cell components grown against an ECM gel (**Figure 1D**). To establish this model, a collagen-I ECM gel was patterned into the middle channel of the OrganoPlate 3-lane Narrow ECM to study different aspects of kidney function and inflammation (**Figure 1A**). The OrganoPlate 3-lane Narrow ECM consists of 40 microfluidic chips embedded in a standard 384-well microtiter plate. Each chip is comprised of three channels that join in the center of the chip: two medium perfusion channels and a gel channel in the middle in which an extracellular matrix (ECM) gel can be patterned through the presence of the phaseguides (**Figure 1B**).^43,44^ The ECM gel compartment in this plate is half the width of the standard OrganoPlate 3-lane 40. This allows more efficient migration of immune cells from the endothelial compartment to the proximal tubule. Upon gelation of the ECM gel, RPTECs were seeded in the top channel against this ECM gel. Next, HUVECs were added to the bottom channel. After several days in culture both cell types formed confluent tubules, after which primary human monocytes were introduced into the lumen of the endothelial vessel (**Figure 1E**). As a result, a renal microfluidic tri-culture model was created including a proximal epithelial tubule (top) and an endothelial vessel with fluorescently labeled monocytes (bottom) as demonstrated in **Figure 1F**. The development of this tri-culture model over time is demonstrated in **supplementary Figure 1**.

### Characterization of the tri-culture model

After establishing the renal tri-culture model, expression of different epithelial and endothelial markers was examined to assess the physiological relevance using an immunofluorescence-based approach in combination with high-content microscopy. A confocal 3D reconstruction of the tri-culture model confirmed the tubular structure and lumen formation of both the RPTEC and HUVEC cultures (**Figure 2A**). Actin and acetylated tubulin, visualizing the cytoskeleton and the primary cilia respectively, were observed in both cell types. This could also be observed at higher magnification max projection images (**Figure 2B,C**). Furthermore, RPTEC and HUVEC tubules demonstrated tight junction formation as shown by immunofluorescence staining of ZO-1 (**Figure 2B,C**) and expression of adherens junction markers VE-cadherin and CD31 in the HUVEC tubule (**Figure 2D,E**). The presence of human primary monocytes was shown by expression of CD45, a cell surface marker of nucleated hematopoietic cells involved in regulation and activation of the immune system (**Figure 2F**).

**Figure 2.**
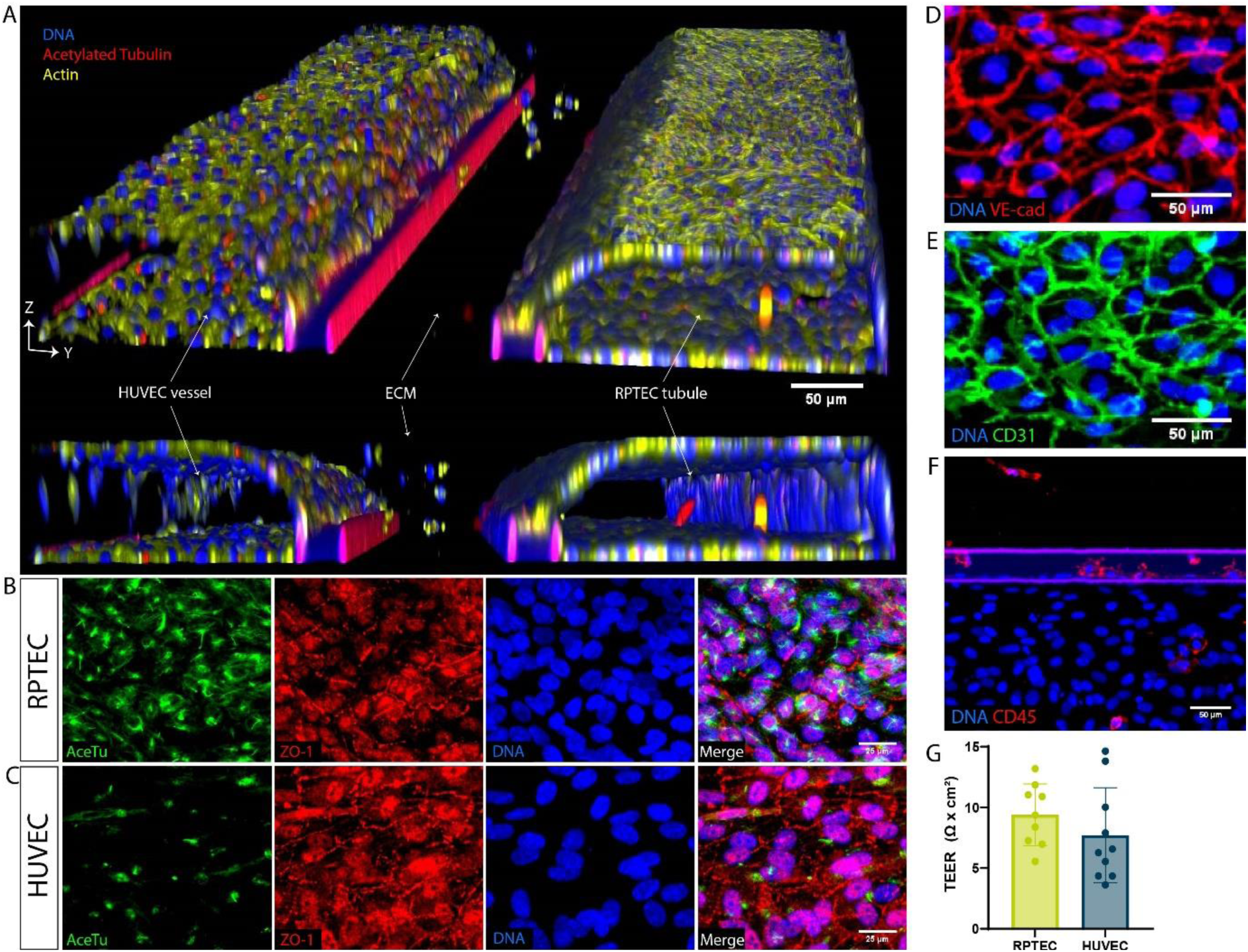
Characterization of the kidney on-a-chip model. (**A**) 3D reconstruction of a confocal z-stack showing both tubular structures with clear lumen grown against a collagen I ECM in the 3-lane Narrow ECM plate at 10X magnification. Cultures were stained for acetylated tubulin (red), actin (yellow), and DNA (blue). (**B,C**) Representative immunofluorescent max projections of the RPTEC or HUVEC cultures respectively, stained for acetylated tubulin (green), zonula occludens I (ZO-1; red), and DNA (blue). (**D**) Max projection of the HUVEC culture stained for VE-cadherin (red) and DNA (blue). (**E**) Max projection of the HUVEC culture stained for CD31 (green) and DNA (blue). (**F**) Max projection indicating part of the HUVEC tubule containing monocytes and ECM compartment with a phaseguide in between. Cultures were stained for CD45 (red) and DNA (blue). (**G**) Epithelial and endothelial barrier function in the model was assessed by measuring TEER at day 6 of culture (n=9-10). All cultures were fixed on day 10 of culture. Scale bars in white = 50 µm (**A,D-F**) or 25 µm (**B,C**).

After observing expression of adherens- and tight junction markers in the renal tri-culture, barrier formation of the epithelial (RPTEC) and endothelial (HUVEC) tubules was examined using TEER (transepithelial/endothelial electrical resistance) measurements. Barrier tightness was measured on day 6 and showed a TEER of 9.4 ± 2.5 and 7.7 ± 3.9 Ω·cm^2^ for RPTEC and HUVEC, respectively.

### Assessing immune-mediated damage

Complement activation as well as recruitment and infiltration of immune cells such as monocytes are key steps of an inflammatory response in the proximal tubule which can lead to significant damage. This complex process involves cellular activation, release of pro-inflammatory cytokines, compromised cell viability, and loss of cells.^45,46^ Here, we examined the immune-mediated effects of renal inflammation in the renal tri-culture model. To this end, renal inflammation was induced using 5% complement activated serum (CAS) which was added to the RPTEC channel and allowed to form a gradient in the chip (**Figure 3A i, ii**). Fluorescently labeled monocytes were added to the lumen of the HUVEC vessel and were tracked in real-time using fluorescent microscopy (**Figure 3A iii, iv**).

**Figure 3.**
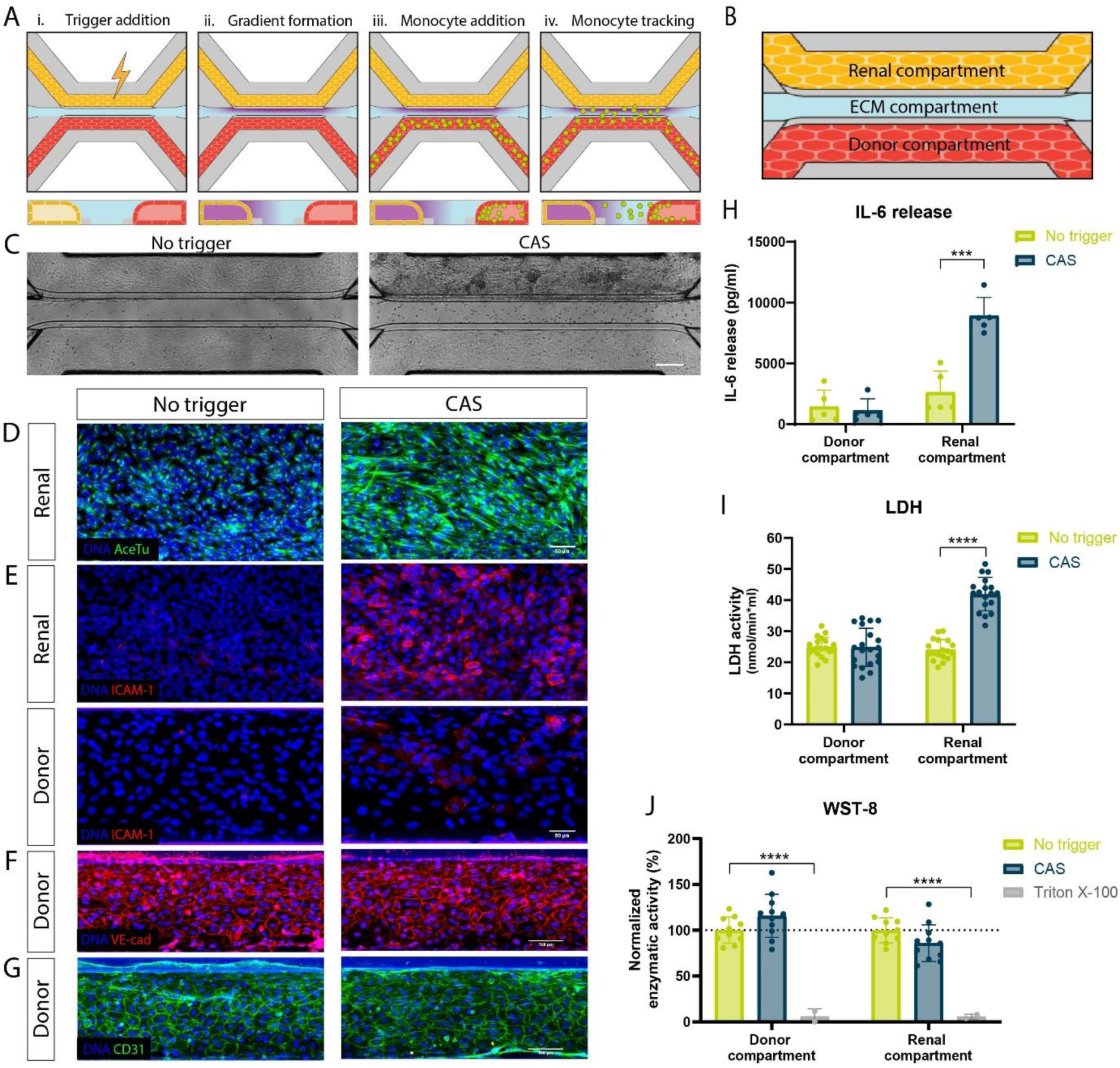
Assessing immune-mediated effects of renal inflammation in the tri-culture model. (**A**) To induce inflammation in the model, a trigger comprised of 5% complement activated serum (CAS) in culture medium was added to the lumen of the RPTEC tubule in the top channel (i) and incubated for 4 hours to establish a gradient (ii). Next, fluorescently labeled monocytes were added to the lumen of the endothelial vessel (iii) and migration was tracked by collecting fluorescent images at different timepoints (iv). (**B**) Schematic representation of the model indicating the nomenclature for each culture channel, namely the donor compartment (bottom perfusion channel), the ECM compartment and the renal compartment (top perfusion channel). (**C**) Phase contrast images of the tri-culture model exposed to no trigger (culture medium) or to 5% CAS for 96h. (**D**) Acetylated tubulin (green) and DNA (blue) immunofluorescent staining in the Renal compartment in non-triggered and CAS-exposed conditions. (**E**) Representative immunofluorescent max projections of the Renal and Donor compartment exposed to no trigger or CAS for 96h and stained for ICAM-1 (red) and DNA (blue). (**F, G**) Max projection of the Donor compartment in non-triggered and CAS-exposed conditions stained for VE-cadherin (red) or CD31 (green) and DNA (blue) (**H**) Secretion of interleukin 6 (IL-6) was assessed in the Renal and Donor compartment in the non-triggered and CAS-exposed conditions at 96h post-exposure. (**I**) Lactate dehydrogenase (LDH) release in supernatant collected from the Donor and Renal compartment after 96h exposure to no trigger or CAS. (**J**) Enzymatic activity of the cultures in the Donor and Renal compartment exposed to no trigger or CAS was determined using WST-8. Triton X-100 was included as a positive control. Graphs showing mean ± standard deviation and individual chips. Data figure D-I monocyte donor 2; data figure J monocyte donor 1. IL-6: n=5, LDH: n=18-20, WST-8: n=2-11. Statistical analysis was performed using multiple unpaired t tests for IL-6 and LDH or two-way ANOVA with Sidak’s multiple comparisons test for WST-8. ***P < 0.001, ****P < 0.0001. All cultures were fixed on day 10 of culture. Scale bars in white = 200 µm (**C**), 50 µm (**D,E**) or 100 µm (**F,G**).

To evaluate the immune-mediated effects on each part of the culture, the chip was divided into the following compartments for analysis: Donor (HUVEC channel), ECM (ECM channel) and Renal (RPTEC channel) (**Figure 3B**). CAS exposure resulted in morphological alterations in the RPTEC channel as demonstrated by cell clustering and slight tubular contraction observed with phase contrast imaging after 96 hours compared to no trigger (**Figure 3C, supplementary Figure 2**). Furthermore, increased monocyte infiltration into the ECM compartment was observed upon exposure to CAS.

Immune-mediated upregulation of both acetylated tubulin, a marker for primary cilia, and ICAM-1 (intercellular adhesion molecule) within the RPTEC tubule were seen as well (**Figure 3D,E**). A slight increase in ICAM-1 expression was observed in the HUVEC vessel upon CAS exposure (**Figure 3E**). Expression of VE-Cadherin and CD31 in the HUVEC vessel (Donor), a marker for adherens junction and vascular adhesion molecule respectively, did not seem to be affected by exposure to CAS (**Figure 3F,G**).

Cell culture supernatant was collected from the Donor and Renal compartment separately after 96h exposure and used to assess the levels of interleukin 6 (IL-6). This cytokine plays a multifaceted role in the immune system and has an influence on pro-inflammatory signaling, recruitment of immune cells, directly impacting epithelial cells, and regulating tubulointerstitial fibrosis.^47^ As shown in **Figure 3H**, no distinction in the detected IL-6 levels was found within the Donor compartment between conditions, but the levels were significantly increased in the Renal compartment after exposure to CAS. IL-6 levels at the start of the assay (0h) were found to be 200-2000 times lower compared to the end of the assay (96h) (**supplementary Figure 3**).

LDH is a cytoplasmic enzyme that is being released into the culture supernatant due to compromised plasma membranes and serves as an indicator for cell damage.^48^ Immune-mediated injury was confirmed by the increased LDH release in the inflamed renal compartment compared to the non-triggered control after 96h exposure (**Figure 3I**). Supernatant collected at the start of the assay (0h timepoint) showed comparable LDH levels between untriggered and triggered conditions (**supplementary Figure 3**). Finally, the metabolic cell activity as an indicator of cell viability was assessed using a WST-8 assay.^49^ No significant differences were observed in the Donor and Renal compartment between inflamed and non-inflamed conditions (**Figure 3J**). Overall, this tri-culture setup with inflammatory trigger provided important insights into the complex processes involved in renal inflammation and allowed us to study different aspects.

### Monocyte migration under healthy and inflamed renal conditions

Monocytes and monocyte-derived cells have distinct roles in the proximal tubule under normal conditions and are actively involved in the immune response during inflammation.^50,51^ Here, we examined the behavior of primary human monocytes in the context of renal inflammation by determining their migratory behavior and compared it to non-inflamed culture conditions.

Primary human monocytes were labeled with a fluorescent Celltracker dye, added to the lumen of the HUVEC vessel and tracked in real-time. The location of the monocytes within the chip was determined at different timepoints and this enabled studying monocyte extravasation and recruitment towards the site of inflammation.

Monocyte migration was examined over time by fluorescent imaging and confirmed elevated migration in the inflamed condition compared to the non-triggered control condition (**Figure 4A**). The development of renal inflammation, monocyte extravasation, and subsequent migration was shown to be a dynamic process as captured in **supplementary Figure 4 (video 1)**. Monocytes were observed to patrol over the vessel wall before extravasating into the ECM, after which monocytes travelled different distances with several eventually reaching the Renal compartment and some even returning to the endothelial tubule. Furthermore, CAS exposure resulted in more, but also faster monocyte migration as observed in both phase contrast and fluorescent imaging.

**Figure 4.**
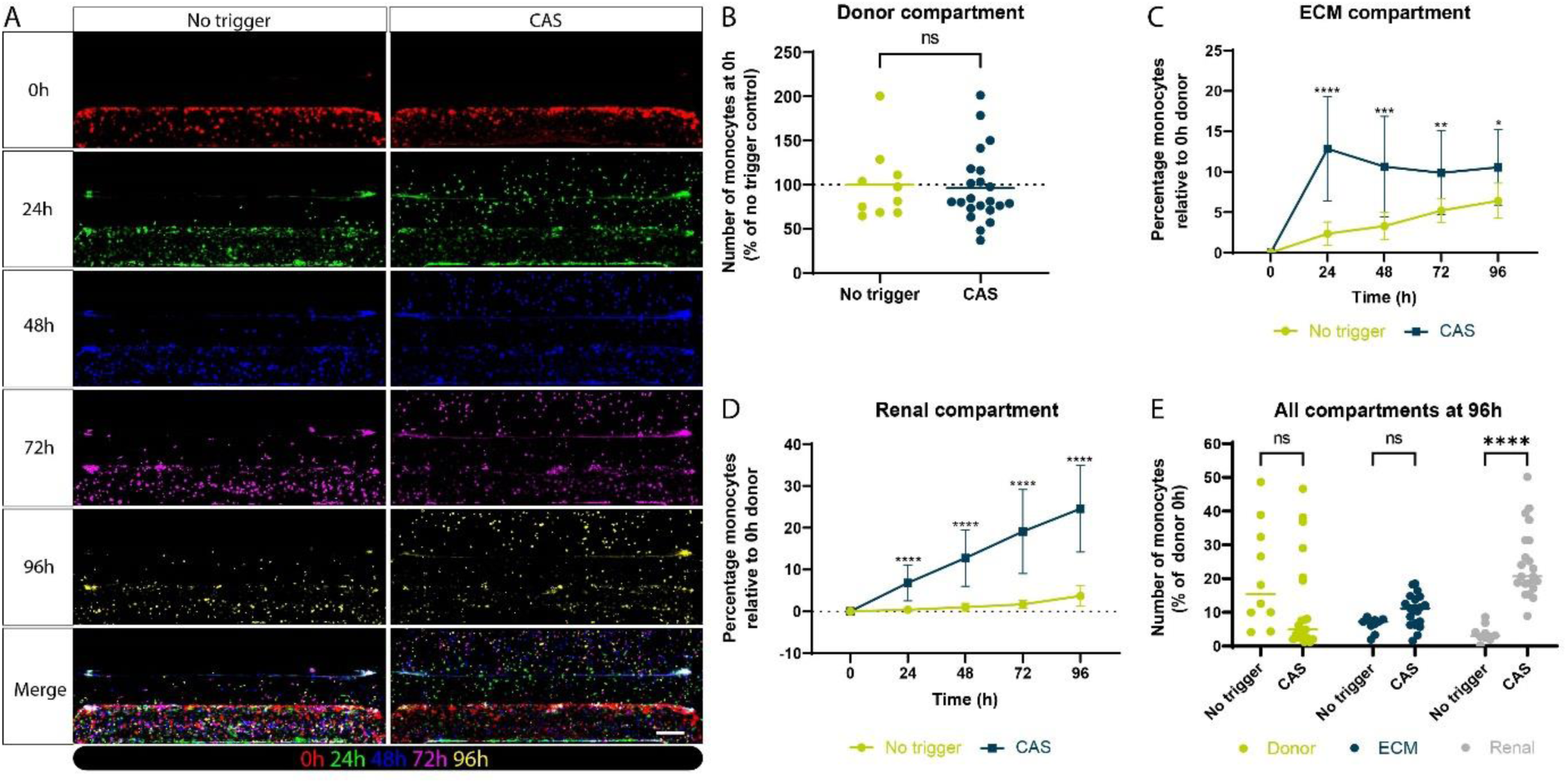
Assessing monocyte migration under healthy or inflamed renal conditions. (**A**) Fluorescent images of the tri-culture model indicating the location of the monocytes at different timepoints in the no trigger or CAS-exposed condition. Monocytes were labeled using Celltracker Orange CMRA (TRITC). To visualize the distance/location of the monocytes within a chip over time, each specific timepoint is displayed with a different color. (**B**) Number of monocytes in the Donor compartment at the start of the assay (0h) normalized to the no trigger control. (**C,D**) Percentage of monocytes that migrated into the ECM or Renal compartment of the chip relative to number of monocytes at start of the assay in the Donor compartment depicted over time comparing no trigger against CAS-exposed condition. (**E**) Percentage of monocytes present in each compartment of the chip (Donor, ECM or Renal) after 96h in the no trigger or CAS-exposed conditions. Graphs showing mean ± standard deviation or individual chips, N=2, n=10-22 chips. Statistical analysis was performed using unpaired t test (B), mixed-effects analysis (C,D), or two-way ANOVA (E); ns = not significant, *P < 0.05, **P < 0.01, ***P < 0.001, ****P < 0.0001. Scale bars in white = 200 µm.

The fluorescent images were used to quantify monocyte adhesion to the endothelial vessel, one of the first steps for monocyte trafficking in response to inflammatory signals. Here, the percentages of monocytes that were present at the start of the assay in the non-triggered and CAS-triggered conditions relative to the no trigger control were determined (**Figure 4B**). Results showed that there were no significant differences between the groups. Next, quantification of the images also allowed us to study monocyte migration by determining the number of cells in each compartment of the chip at different timepoints. The percentage of monocytes in the ECM and Renal compartment over time in the non-triggered and CAS-triggered conditions relative to the number of monocytes at the start of the assay in the Donor compartment are depicted in **Figure 4C and 4D**. Exposure to CAS significantly increased monocyte migration into both compartments at all timepoints between 24-96 hours compared to the non-triggered condition. Upon exposure to CAS, there is a steep significant increase of monocytes migrating into the ECM compartment after 24 hours which stabilizes over time or even slightly decreases (**Figure 4C**). This is likely linked to the observed linear increased monocyte migration into the Renal compartment (**Figure 4D**). The distribution of the monocytes in the chip after 96-hour exposure to no trigger or CAS showed a significant difference in the Renal compartment (**Figure 4E**).

### Assessing donor variability in the renal inflammation model

To determine the robustness and reliability of the renal inflammation model and its applicability for personalized medicine and drug development, it is crucial to understand the effect of donor differences. To examine donor variability, primary human monocytes from 4 different donors were used to establish the renal tri-culture model and were challenged with CAS to induce inflammation. Monocyte adhesion and migration were assessed and quantified using fluorescent imaging over time.

Monocyte adhesion at the start of the assay showed no significant differences between the no trigger and CAS-exposed conditions for all donors (**Figure 5A**). However, the variability within each donor is different where donor 1 showed the highest variability and donor 4 the least. Next, the percentage of monocytes in each compartment was determined relative to the number of monocytes in the Donor compartment at the start of the assay (0h). **Figures 5B-D** show the results of the 96-hour timepoint in the Donor, ECM, and Renal compartment, respectively. At this timepoint, clear differences between donors could be observed in each of the compartments. Donor 3 revealed increased presence of monocytes in the Donor compartment compared to the other donors. Upon exposure to CAS, a decreased trend could be observed for all donors, and was significantly different for donor 3 (**Figure 5B**). In the ECM compartment, an increased percentage of monocytes was observed upon exposure to CAS for all donors, and this was significantly increased for donors 2 and 3. Finally, the effect of CAS was homogeneously observed in the Renal compartment as shown by significantly elevated migration for all four donors. The biggest difference between no trigger and CAS was observed for donor 4. An overview with data of the other timepoints of donor 1, 3 and 4 can be found in **supplementary Figure 5**. These graphs revealed further differences in dynamics of monocyte adhesion and migration over time between donors.

**Figure 5.**
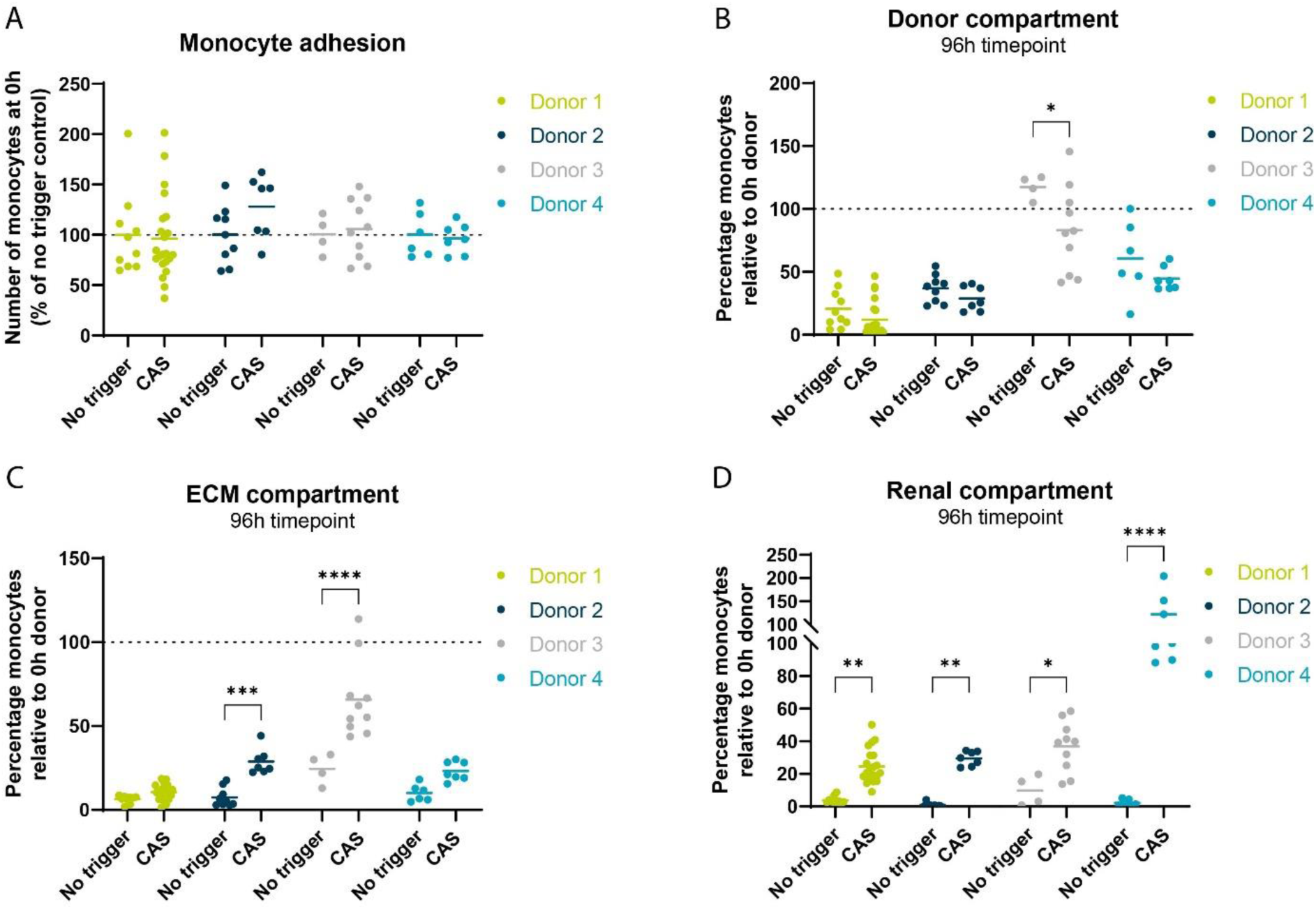
Evaluating migration and adhesion of different monocyte donors. (**A**) Number of monocytes in the Donor compartment at the start of the assay (0h) normalized to the no trigger control separately for each monocyte donor. (**B-D**) Percentage of monocytes present in each compartment of the chip (Donor, ECM or Renal) after 96h exposure to no trigger or to 5% CAS represented for 4 different monocyte donors. Graphs showing mean and individual chips. Donor 1: N=2, n-10-22; Donor 2: N=1, n=7-9; Donor 3: N=1, n=4-10; Donor 4: N=2, n=6-7. Statistical analysis was performed using two-way ANOVA; *P < 0.05, **P < 0.01, ***P < 0.001, ****P < 0.0001

### Effect of immune modulatory compounds on monocyte migration

The renal inflammation assay demonstrated to be an effective tool to determine monocyte behavior under healthy and inflamed conditions. Therefore, the applicability of the assay to evaluate effects of immune modulatory compounds on monocyte migration was tested. Two compounds, namely compound A and B were evaluated in the presence of 5% CAS where compound A targets the inflammatory trigger and compound B targets the monocytes directly.

Based on the compound characteristics, different exposure strategies were applied. Different concentrations of compound A were tested, and an optimal concentration was selected for further experiments (data not shown). Here, the effect of adding compound A to the Renal compartment, Donor compartment, or to both compartments was determined by using monocyte donor 4. Compound B was added to the Renal compartment and tested at three different concentrations using monocyte donor 1.

Both compounds showed significant inhibition of monocyte migration displayed in **Figure 6B and 6D** by the decreased percentage of monocytes reaching the Renal compartment when exposed to both CAS and compound A or B. Results showed that all the treatment administration methods tested for compound A significantly inhibited monocyte migration into the Renal compartment after 96h exposure (**Figure 6B**). Also, migration into the ECM compartment was significantly inhibited upon exposure to all compound A treatment methods as compared to CAS at 96h (**Figure 6A**). For compound B, all three tested concentrations resulted in equivalent migration inhibition into the Renal compartment (**Figure 6D**). On the contrary, monocyte migration into the ECM compartment was not affected by compound B (**Figure 6C**). Results of the Donor, ECM, and Renal compartment over time can be found in **supplementary Figure 6**. In summary, these results showed that treatment with immune modulatory compounds can diminish monocyte migration in the light of renal inflammation in the tri-culture model.

**Figure 6.**
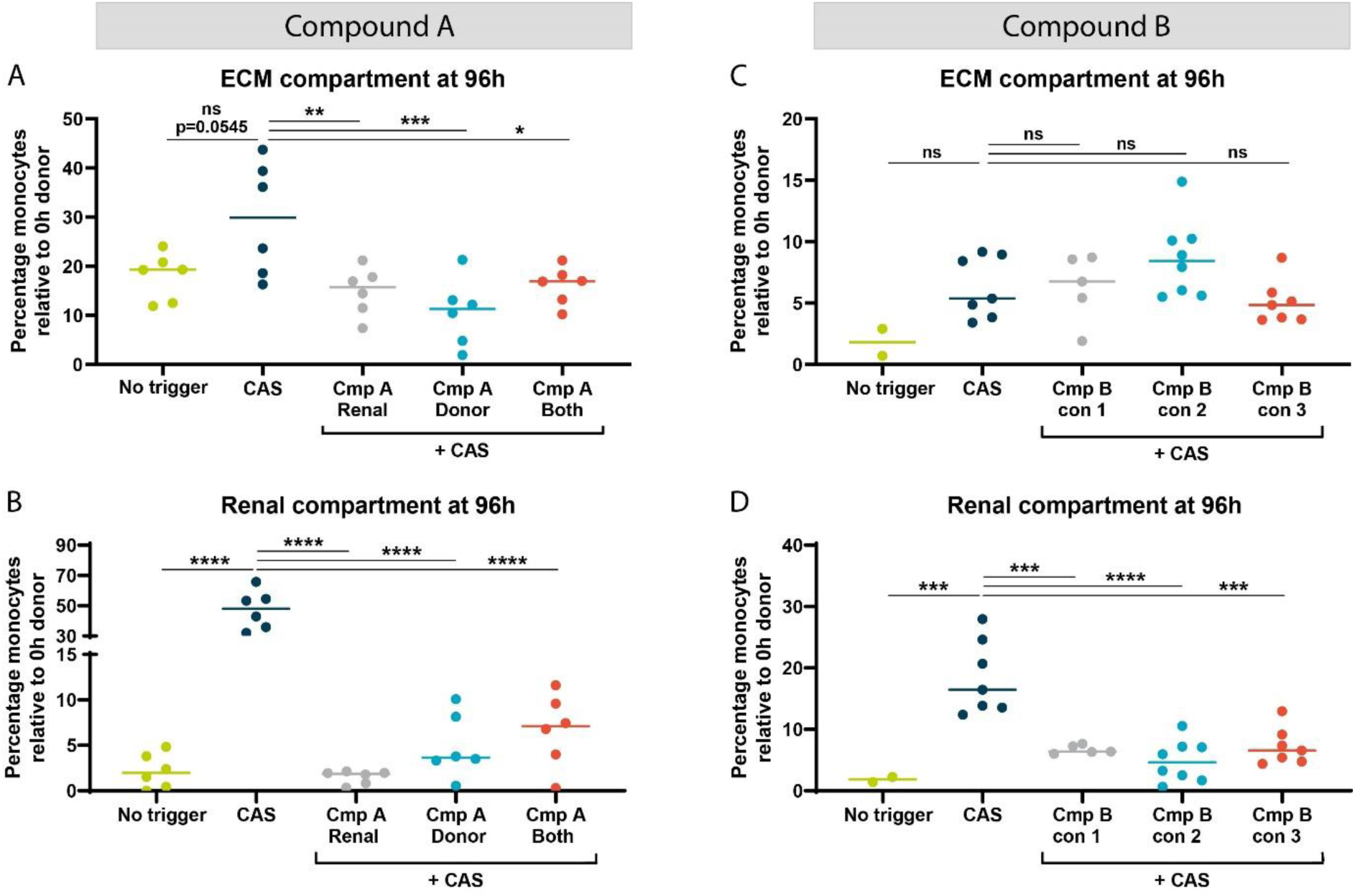
Effect of immune modulatory compounds on monocyte migration. Compound A was added to the Renal compartment, Donor compartment or to both compartments of the chip and tested using monocyte donor 4. Compound B was tested at 3 different concentrations (#1, 2 or 3), added to the Renal compartment, and tested using monocyte donor 1. (**A,C**) Percentage of monocytes present in the ECM compartment after exposure to no trigger, CAS, or CAS + Compound A or B respectively after 96h. (**B,D**) Percentage of monocytes present in the Renal compartment after exposure to no trigger, CAS, or CAS + Compound A or B after 96h. Graphs showing mean and individual chips. Compound A: N=1, n=5-6; Compound B: N=1, n=2-8. Statistical analysis was performed using one-way ANOVA; ns = not significant, *P<0.05, **P < 0.01, ***P < 0.001, ****P < 0.0001.

## Discussion

Numerous renal diseases are recognized to be driven by the immune system, where dysregulated immune responses are key contributors to their pathogenesis. Despite this recognition, the development of targeted therapies has been challenging as knowledge of the underlying mechanism and complex interactions remains insufficient. Recent advancements in the field offer promising avenues for exploring the interplay between renal cells and immune cells and their role in the development of renal inflammation and diseases. This study describes the establishment of a human immunocompetent 3D *in vitro* co-culture model of the proximal tubule in a high-throughput microfluidic platform that can be used to study renal functionality and inflammatory processes. The model is especially valuable for its ability to incorporate monocytes, an understudied cell type that is becoming increasingly relevant to renal diseases in CKD and beyond.^52–54^

The model incorporated RPTEC in the top compartment and HUVECs in the bottom compartment cultured under flow and in direct contact with a collagen-I ECM gel resulting in the formation of polarized tubular structures. As an immune component, human primary monocytes of different donors were added to the lumen of the endothelium. Renal inflammation was successfully induced using complement activated serum as evident by epithelial morphological changes, increased expression of adhesion molecules, release of pro-inflammatory cytokines, and reduced epithelial viability. Realtime migratory behavior of monocytes showed increased extravasation and migration towards the ECM and Renal compartment upon exposure to CAS with donor-to-donor differences observed. Finally, immune modulatory compounds showed efficacious inhibition of monocyte migration under inflammatory conditions in the microfluidic co-culture model.

The microfluidic platform used in this study is a custom-made prototype plate utilizing the same format as the OrganoPlate 3-lane 40 platform but with different channel dimensions including a smaller ECM channel. Accordingly, it facilitated enhanced migration of monocytes from the endothelium towards the epithelium within a set timeframe (data not shown). Compared to migration studies in Transwell systems, a microfluidic platform allowed for better control of the microenvironment, flexibility to adjust experimental complexity, improved visualization and analysis, and higher throughput.^55–57^ For example, the flow rate and concentration gradients could be precisely controlled and adjusted to recapitulate dynamic environmental cues. Furthermore, perfusion flow was generated without the use of pumps by placing the OrganoPlate on an interval rocker, generating gravity-driven fluid flow in a bidirectional manner. With the used rocker settings, a peak shear stress of 1.2 dyne/cm^2^ was established in both channels.^58^ This falls within the *in vivo* estimated range of 0.3 to 1.2 dyne/cm^2^ for the human proximal tubule but is lower compared to the reported range of 4 to 12 dyne/cm^2^ for endothelial cells.^59,60^ Even though the flow is bidirectional, fluid flow is an important cue for proximal tubule cells and has been shown to increase polarization, transporter functionality, and morphology.^61–63^ Besides fluid shear stress, direct cell-ECM interaction also contributes to improved proximal tubule characteristics,^64^ which is possible in the used microfluidic platform as there is no artificial membrane. In line with these reports, we showed polarized 3D tubular structures with a clear lumen and expression of actin and acetylated tubulin indicating the presence of the cytoskeleton and primary cilia.

Barrier formation to control the passage of molecules through the cell layer is another crucial characteristic of the proximal tubule.^65,66^ In our model, barrier formation was shown by expression of tight and adherens junction markers ZO-1, in both RPTEC and HUVEC, and VE-cadherin and CD31 in the HUVEC. Tightness of the barrier was assessed using TEER measurements and revealed an average TEER of 9.4 ± 2.5 Ω·cm^2^ for the RPTEC which is in line with the reported *in vivo* range of 6.6 to 11.6 Ω·cm^2^.^67,68^ For HUVECS, an average TEER of 7.7 ± 3.9 Ω·cm^2^ was observed. Direct *in vivo* TEER measurements of HUVECs or blood vessels in general are lacking. However, *in vitro* HUVEC TEER values have been reported to range from 10 to 1000 Ω·cm^2^ and vary based on the experimental conditions and environmental factors.^69,70^ Although beyond the scope of this study, it would be interesting to study the barrier function in real-time under inflammatory conditions through timelapse TEER measurements as shown by Ehlers *et al.* for vascular inflammation.^71^

Key pathogenic factors of a pro-inflammatory response in renal tubules are complement activation and immune cell infiltration both driving tubulointerstitial damage.^72,73^ To induce renal inflammation in the model, complement activated serum was added to the Renal compartment and allowed to form a gradient in the chip. Morphological alterations in the RPTEC tubules were observed upon exposure to CAS. Renal tubular epithelial cells can produce but also activate complement via the alternative pathway.^74,75^ In turn, the tubular epithelium becomes injured and/or activated resulting in the synthesis and release of pro-inflammatory cytokines, including TNFα and IL-6, release of reactive oxygen species, and increased synthesis of matrix proteins.^76^ In line with these reports, we observed significantly increased IL-6 release from the Renal compartment as well as increased ICAM-1 and acetylated tubulin expression in RPTECs in the presence of CAS. Increased expression of ICAM-1 on renal tubular epithelial cells under inflammatory conditions has been reported and can potentially serve as a biomarker to predict disease progression.^77–79^ ICAM-1 can be expressed on both the apical and basolateral side of the proximal epithelium and has an important role in regulating the response to infiltrating immune cells.^80^ In addition to increased expression in RPTECs, elevated ICAM-1 expression was also observed in the HUVEC tubule in accordance with literature indicating its role in regulating transendothelial migration in response to inflammation.^81^

Acetylated tubulin is found in primary cilia, but also in the centrioles and flagella of epithelial cells, and is regulating cell polarization, proliferation, development, and migration.^82^ Dysregulated tubulin acetylation has been associated with tubular cell dysfunction, and if not treated can advance to tubular apoptosis and necrosis.^83^ The increased expression and altered appearance of acetylated tubulin observed in the RPTEC tubule indicate early signs of tubular damage.

Finally, viability of the cultures was determined by detecting released LDH and measuring enzymatic cell activity using WST-8. Cells undergoing apoptosis, necrosis or other forms of damage will release LDH from the cytoplasm into the supernatant as a signal of impaired cell membranes.^84^ Under inflammatory conditions, we observed significantly increased LDH in the Renal compartment suggesting damaged RPTECs. This is not the case for the Donor compartment containing the HUVEC culture. On the contrary, no clear effect upon CAS exposure was detected in the WST-8 assay. A possible explanation for this might be that epithelial cells undergoing programmed cell death maintain metabolic activity during certain stages of the process. Sustained metabolic activity during early stages of apoptosis has been crucial for ensuring proper regulation of the controlled breakdown of cellular components.^85^ To further investigate immune-mediated effects on the culture viability and to improve our understanding of the type of cell death involved, it would be interesting to prolong the experiment to check the viability at later stages and include other readouts to determine caspase activity and DNA fragmentation.^86^

During complement activation, the anaphylatoxins C3a and C5a are produced and establish a gradient primarily aimed to induce chemotaxis.^73^ These and other complement proteins activate resident and infiltrating cells leading to production and secretion of pro-inflammatory cytokines that further stimulate leukocyte recruitment and infiltration.^87^ One of the first responders recruited to the inflammatory tubulointerstitial microenvironment are monocytes, which adhere to activated endothelium and transmigrate into the interstitial space. To study monocyte behavior in normal and inflammatory conditions, primary human monocytes were fluorescently labeled and perfused through the lumen of the HUVEC tubule. With the established workflow in our setup, we were able to visually detect and quantify the number, location, and movement of monocytes in real-time. No differences in monocyte adhesion to the endothelial vessel wall at the start of the assay was observed between the triggered and non-triggered group. This was observed with each of the four monocyte donors. One explanation could be that endothelial activation occurs gradually upon exposure to complement proteins and requires additional time for the synthesis and expression of adhesion molecules.^88^ On the contrary, significant increased monocyte migration was observed in the triggered condition towards the ECM and Renal compartment. Overall, the impact of CAS was consistently observed across all tested monocyte donors, indicating its efficacy as a robust trigger for evaluating monocyte migration in the context of renal inflammation.

To examine the robustness of the assay, it is crucial to understand the effect of donor differences. Hence, four monocyte donors were tested and differences in the number of adhering monocytes to the endothelial vessel wall were observed, both at the beginning of the assay and over time. Moreover, CAS proved to be a potent inducer of migration; however, the extent of migration and location of the monocytes varied among donors. Monocytes can be classified in three major subpopulations, namely, classical, intermediate, and non-classical. Distinguishment is mainly based on the expression of CD14 and CD16 on the surface and their function in homeostasis or disease including inflammatory response and migratory potential.^89^ This heterogeneity exists between the subsets but has also been demonstrated between healthy and disease conditions and between individuals.^89^ This could explain the observed differences in adhesion and migration between monocyte donors in our assay. Understanding the role of each of the subtypes in the context of renal inflammation, may provide new avenues for targeted therapies and personalized medicine. In an initial test, we examined the migratory behavior of isolated monocyte subsets in triggered and non-triggered conditions (data not shown). Although shown to be feasible, more time and effort is required to obtain an optimized workflow. Additionally, further characterization of the morphology of the migrated monocytes into the ECM channel could provide valuable information on the subsets as well as the differentiation towards macrophages and the role of subtypes in renal inflammation.^90^

The imaging compatibility of the OrganoPlate enabled us to not only track and quantify monocyte migration but also to examine morphological alterations in the epithelial and endothelial tubules and their interaction with monocytes over time. As shown in supplementary figure 4, timelapse videos were captured of the triggered and non-triggered cultures and revealed monocyte patrolling behavior across the endothelial wall and differences in movements towards the Renal compartment. Using advanced data analysis software or AI provides a potential next step to support our knowledge of interactions in this complex microenvironment and detect abnormalities associated with disease conditions.

To provide initial proof-of-concept data that this model can be used to assess investigational therapeutics and serve as a preclinical predictive tool, the cultures were exposed to two immune modulatory compounds under inflammatory conditions. Targeting different components of the inflammatory environment, either the trigger or the monocytes, both compounds showed significant reduction in monocyte migration into the Renal compartment. In addition to assessing the number of migrated monocytes, this assay would also enable detection of the tendency of migration, such as speed, distance, optimal time window, and location. Interestingly, the effect of both compounds varies within the ECM channel, where compound A significantly decreased monocyte migration and compound B did not exhibit the same effect. The level of migration between the no trigger and CAS conditions varies between the compounds as different monocyte donors were utilized. To fully validate the effect of the compounds, it would be advisable to evaluate them on multiple monocyte donors. This could also provide valuable information for patient stratification in later stages of drug development. Using patient-derived monocytes in the assay would enable the assessment of migratory capacity within a renal inflammatory environment and facilitate personalized testing of various (novel) therapeutics.

Crosstalk between the proximal tubule epithelium and peritubular capillaries is crucial for the exchange of signals in the tubulointerstitium and has been shown to influence renal function both in health and disease.^91^ Although the distance in our model exceeds the approximate distance of the *in vivo* human tubular interstitium^92^, which generally does not surpass the diameter of a single cell, it is smaller compared to other microphysiological co-culture models.^93–95^ Additionally, the current size of the tubulointerstitial space in the Narrow ECM plate, represented by the ECM channel, facilitated accurate monocyte quantification and enhances visual interpretability. To study the tubulo-vascular crosstalk, secretion of soluble factors can be determined in our model as shown by the release of IL-6 and LDH. Further investigation of these factors with omics or ELISA analysis can advance our understanding of the molecules secreted and altered under specific (disease) conditions, potentially revealing novel targets for drug development, or identifying new biomarkers.

The versatility of the assay in modulating the system and controlling various environmental factors including cell ratios, stimulus, exposure strategies, and real-time imaging represents significant advantages which cannot be achieved with animal models. By exchanging the disease trigger, the model could be applied for studying other specific renal diseases related to AKI or CKD, such as ischemia-reperfusion injury. Finally, the complexity of the model can be tailored and further increased by incorporation of fibroblasts or other immune cells, such as T cells (**supplementary Figure 7**) or neutrophils.

In conclusion, we successfully established a human immunocompetent 3D *in vitro* co-culture model comprising renal proximal tubule epithelial cells, endothelial cells, and perfused monocytes in a high-throughput microfluidic platform. We were able to induce renal inflammation using complement activated serum and detect immune-mediated damage and associated monocyte migratory behavior. Finally, we demonstrated the use of the assay for assessing effects of immune modulatory compounds in the context of renal inflammation. The model can be applied for fundamental research on renal functionality in health and disease but also for drug screening due to the platform’s compatibility with automation and relatively high throughput. Overall, the described proximal tubule model has high potential to fill the gap that currently exists to study renal inflammation preclinically.

## Supporting information

Supplementary document_Renal inflammation

## Authorship contributions

**Linda Gijzen:** conceptualization, methodology, formal analysis, investigation, writing – original draft, supervision, project administration **Marleen Bokkers:** methodology, formal analysis, investigation, writing – review and editing **Richa Hanamsagar:** methodology, resources **Thomas Olivier:** software **Todd Burton:** resources **Laura Tool:** formal analysis, investigation **Mouly Rahman:** methodology, resources, writing – review and editing **John Lowman:** conceptualization, methodology, project administration **Virginia Savova:** conceptualization, methodology, resources, writing – review and editing, supervision **Terry Means:** conceptualization, methodology, resources, writing – review and editing, supervision **Henriette Lanz:** conceptualization, writing – review and editing, supervision

## Acknowledgements

We would like to thank Frederik Schavemaker for generating artist impressions of the model. We would like to extend our gratitude to Shinji Kasahara, David Habiel and Vladimir Litvak for their contributions and input to this work.

