## Supplementary document_Renal inflammation for "An immunocompetent human kidney on-a-chip model to study renal inflammation and immune-mediated injury"

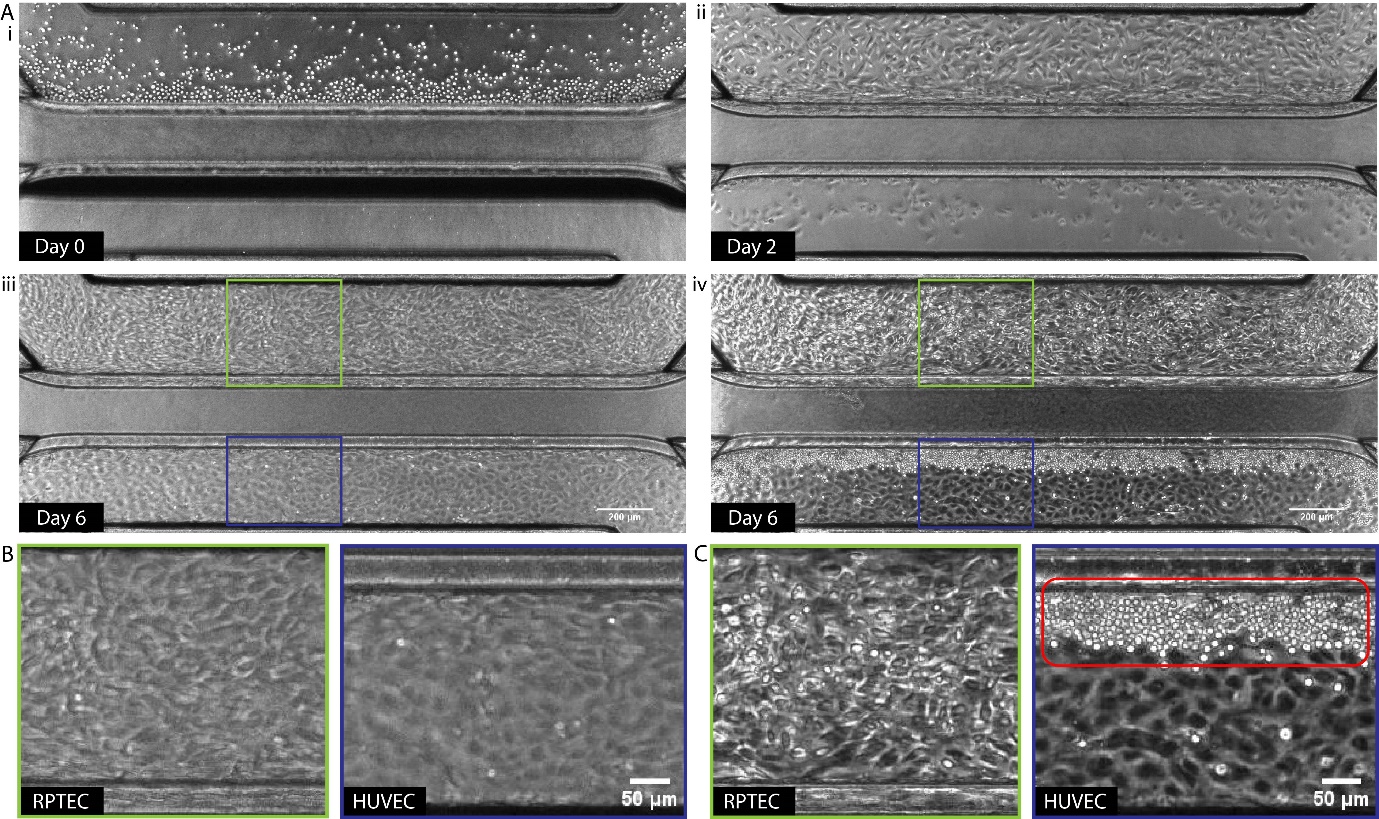

**Supplementary figure 1.** *Development of the tri-culture model over time.* (**A**) Phase contrast images of the cultures in the OrganoPlate at different timepoints. After the ECM gel has been loaded and incubated overnight, human RPTECs were seeded in the top perfusion channel against the ECM gel on Day 0 of the culture (i). On Day 2, HUVECs were seeded in the bottom perfusion channel (ii). Cultures formed confluent tubular structures on Day 6 of culture (iii) after which monocytes were seeded into the lumen of the HUVEC vessel (iv). (**B,C**) Zoom-in images of the RPTEC or HUVEC tubule at Day 6 of culture before and after monocyte addition, respectively. Monocytes were clearly visible in the lumen of the HUVEC vessel lining the ECM-HUVEC interface (red rectangle). Scale bars in white = 200 µm (**A**) or 50 µm (**B,C**).

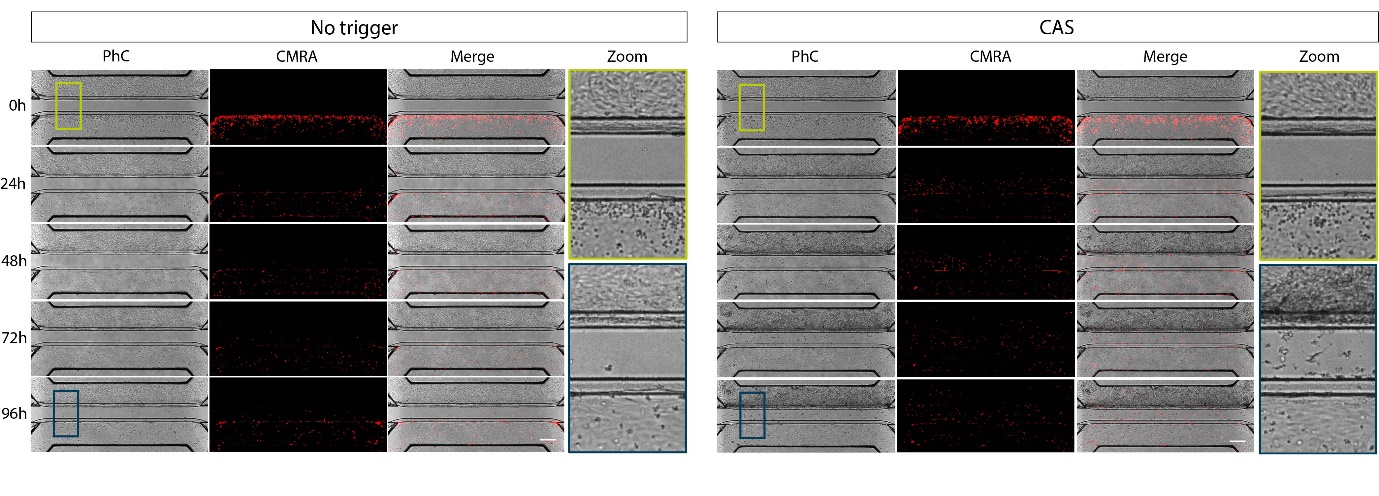
**Supplementary figure 2.** *Morphology assessment and monocyte migration over time.* Phase contrast and fluorescent images of the tri-culture model in the no trigger or CAS-exposed conditions at 0-96 hours. Monocytes were labeled using Celltracker CMRA dye which allow tracking of their location inside the chip. Zoom-in phase contrast images indicated morphological alterations or presence of monocytes at the start of the assay (0h; green marked) and at the end (96h; blue marked). Scale bars in white = 100 µm.

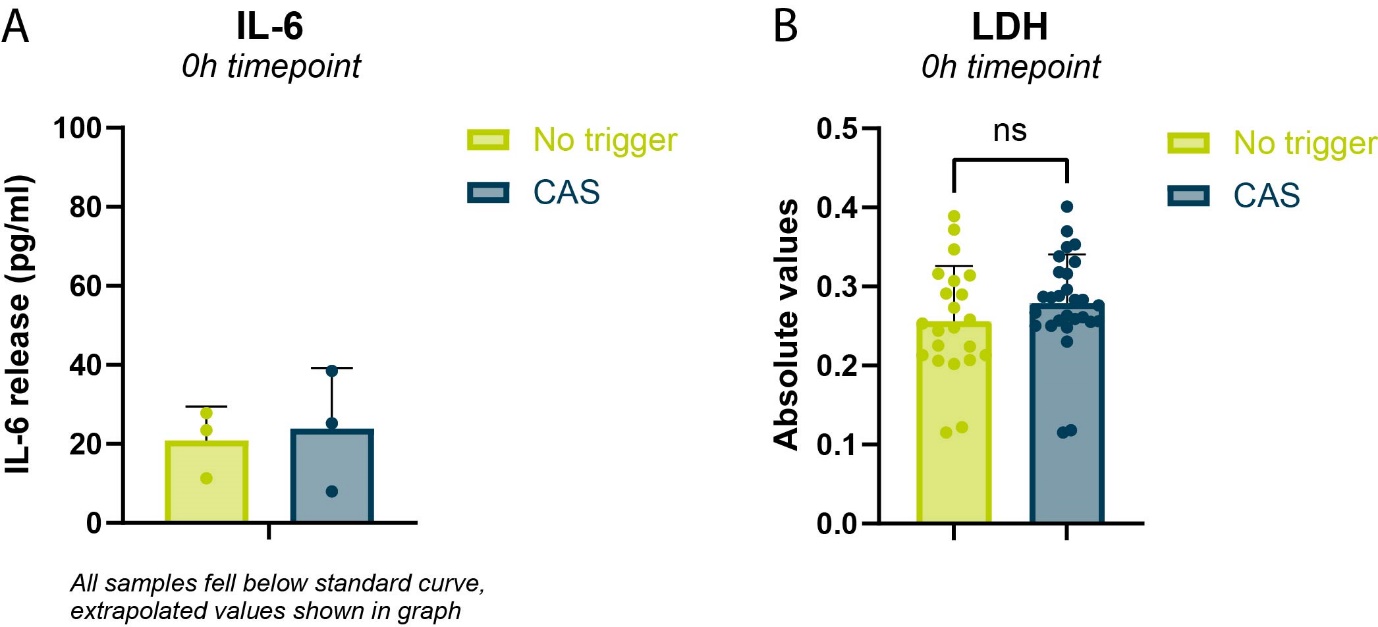

**Supplementary figure 3.** *Assessment of baseline IL-6 and LDH release at start of the assay.* (**A**) Secretion of interleukin 6 (IL-6) was assessed in the non-triggered and CAS-exposed conditions at the start of the assay (0h timepoint). All samples fell below the standard curve and the shown values were extrapolated. (**B**) Lactate dehydrogenase (LDH) release in supernatant of the no trigger or CAS-exposed cultures at the start of the assay (0h timepoint). Graphs showing mean ± standard deviation. Figure A: n=3, Figure B: n=22-28. Statistical analysis was performed using an unpaired t-test for LDH (fig B).

See supplementary figure 4 – video 1

**Supplementary figure 4.** *Real time effect of CAS on morphology and monocyte migration in the kidney on-a-chip model.* Timelapse imaging was performed by capturing phase contrast (**A,B**) and fluorescent (**C,D**) images of the tri-culture model exposed to no trigger or CAS. Every 20 min an image was taken for up to 96 hours. Monocytes were labeled with CMRA Celltracker dye which allowed tracking their location inside the chip. Timelapse video = 5fps.

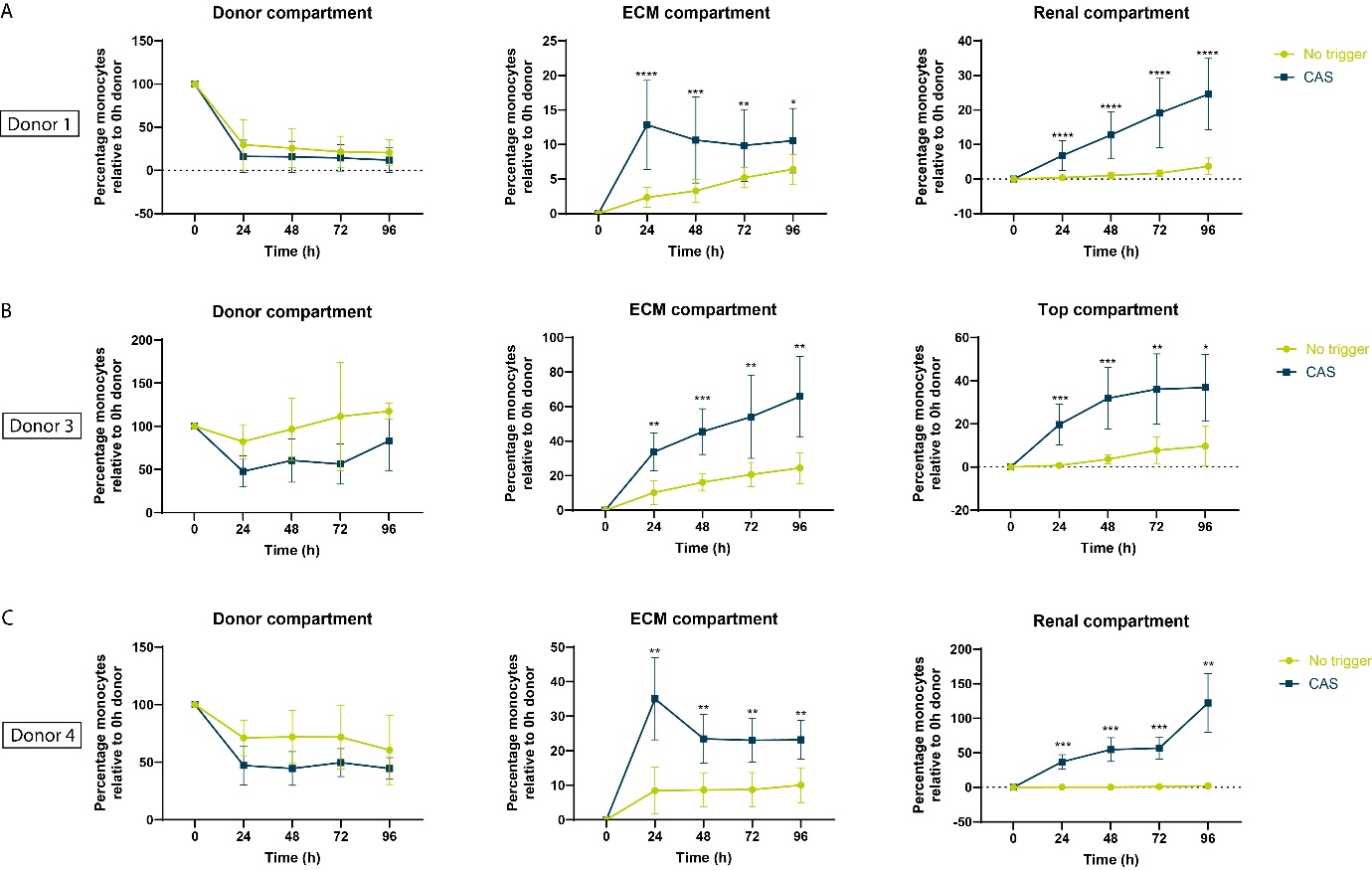

**Supplementary figure 5.** *Migration and adhesion of different monocyte donors.* (**A-C**) Percentage monocytes present in each compartment of the chip (Donor, ECM or Renal) at 0, 24, 48, 72 and 96h after exposure to no trigger or to CAS for Donor 1 (**A**), Donor 3 (**B**) and Donor 4 (**C**). Graphs showing mean ± standard deviation. Donor 1: N=2, n-10-22; Donor 2: N=1, n=7-9; Donor 3: N=1, n=4-10; Donor 4: N=2, n=6-7. Statistical analysis was performed using mixed-effects analysis; *P < 0.05, **P < 0.01, ***P < 0.001, ****P < 0.0001.

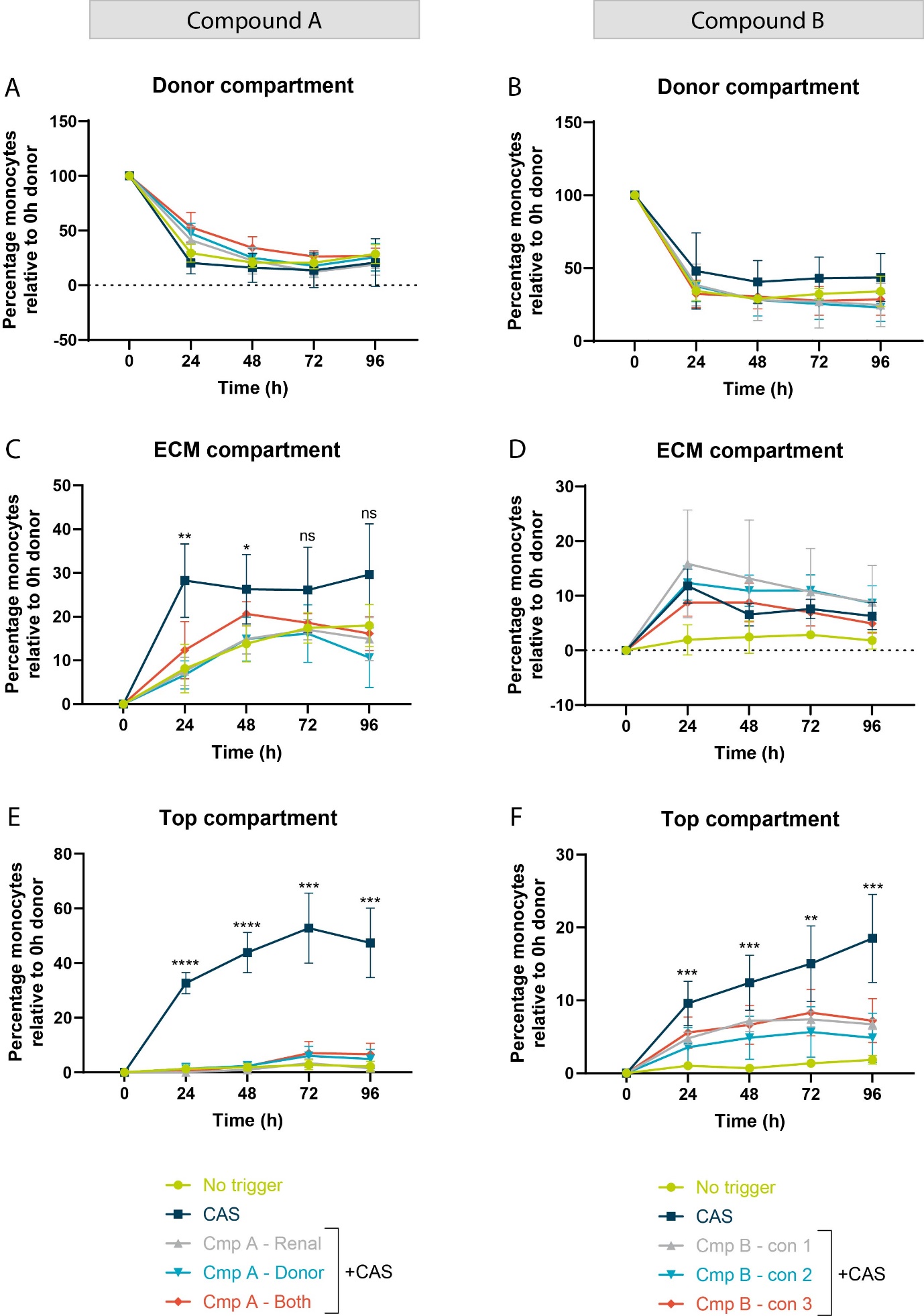

**Supplementary figure 6.** *Effect of immune modulatory compounds on monocyte adhesion and migration.* Compound A was added to the Renal compartment, Donor compartment or to both compartments of the chip and tested with monocyte donor 4. Compound B was tested at 3 different concentrations (#1, 2 or 3) and added to the Renal compartment and tested with monocyte donor 1. Percentage of monocytes present in the Donor (**A,B**), ECM (**C,D**) and Renal (**E,F**) compartment after exposure to no trigger, CAS, or CAS + Compound A or B over time. Graphs showing mean ± standard deviation, Cmp A: N=1, n=5-6; Cmp B: N=1, n=2-8. Statistical analysis was performed using two-way ANOVA; ns = not significant, *P<0.05, **P < 0.01, ***P < 0.001, ****P < 0.0001.

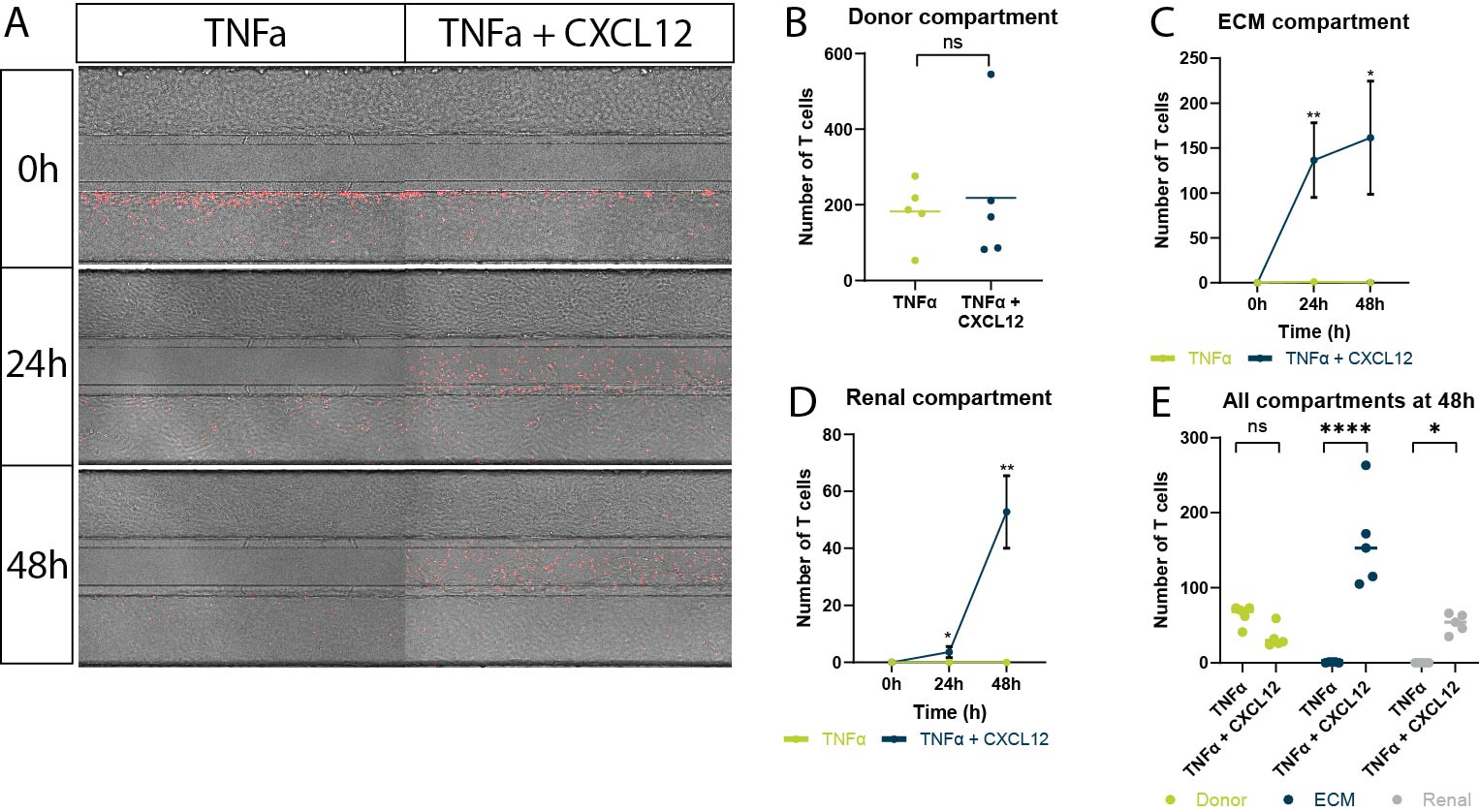

**Supplementary figure 7.** *CD8+ memory T cell migration in the proximal tubule model.* (**A**) Phase contrast and fluorescent overlay images of the tri-culture model indicating the location of the T cells in red at 0, 24 and 48h in the TNFα or TNFα + CXCL12 condition. CD8+ memory T cells were labeled with a fluorescent tracker dye (Cy5). TNFα (2.25 ng/mL) was added to the HUVEC compartment 4 hours before T cell addition. CXCL12 (800 ng/mL) was added to the Renal compartment at the same time as addition of the T cells to the HUVEC compartment. (**B**) Number of T cells in the Donor compartment at the start of the assay (0h). (**C-D**) Number of T cells over time in the ECM and Renal compartment respectively. (**E**) Number of T cells present in each compartment (Donor, ECM or Renal) after 48h in the TNFα +/- CXCL12 exposed conditions. Graphs showing mean ± standard deviation or individual chips, N=2, n=5 chips. Statistical analysis was performed using unpaired t test (B), or two-way ANOVA (C-E); ns = not significant, *P < 0.05, **P < 0.01, ****P < 0.0001.
